# The Metabolic Usage and Glycan Destinations of GlcNAz in *E. coli*

**DOI:** 10.1101/2023.08.17.553294

**Authors:** Alexander Eddenden, Manoj K. Dooda, Zachary A. Morrison, Adithya Shankara Subramanian, P. Lynne Howell, Jerry M. Troutman, Mark Nitz

## Abstract

Bacteria use a diverse range of carbohydrates to generate a profusion of glycans, with amino sugars such as *N*-acetylglucosamine (GlcNAc) being prevalent in the cell wall and in many exopolysaccharides. The primary substrate for GlcNAc-containing glycans, UDP-GlcNAc, is the product of the bacterial hexosamine pathway, and a key target for bacterial metabolic glycan engineering. Using the strategy of expressing NahK, to circumvent the hexosamine pathway, it is possible to directly feed the analogue of GlcNAc, *N*-azidoacetylglucosamine (GlcNAz), for metabolic labelling in *E. coli*. The cytosolic production of UDP-GlcNAz was confirmed using fluorescence assisted polyacrylamide gel electrophoresis. The key question of where GlcNAz is incorporated, was interrogated by analyzing potential sites including: peptidoglycan (PGN), the biofilm-related exopolysaccharide poly-β-1,6-*N*-acetylglucosamine (PNAG), lipopolysaccharide (LPS) and the enterobacterial common antigen (ECA). The highest levels of incorporation were observed in PGN with lower levels in PNAG and no observable incorporation in LPS or ECA. The promiscuity of the PNAG synthase (PgaCD) towards UDP-GlcNAz *in vitro* and lack of undecaprenyl-pyrophosphoryl-GlcNAz intermediates generated *in vivo* confirmed the incorporation preferences. The results of this work will guide the future development of carbohydrate-based probes and metabolic engineering strategies.

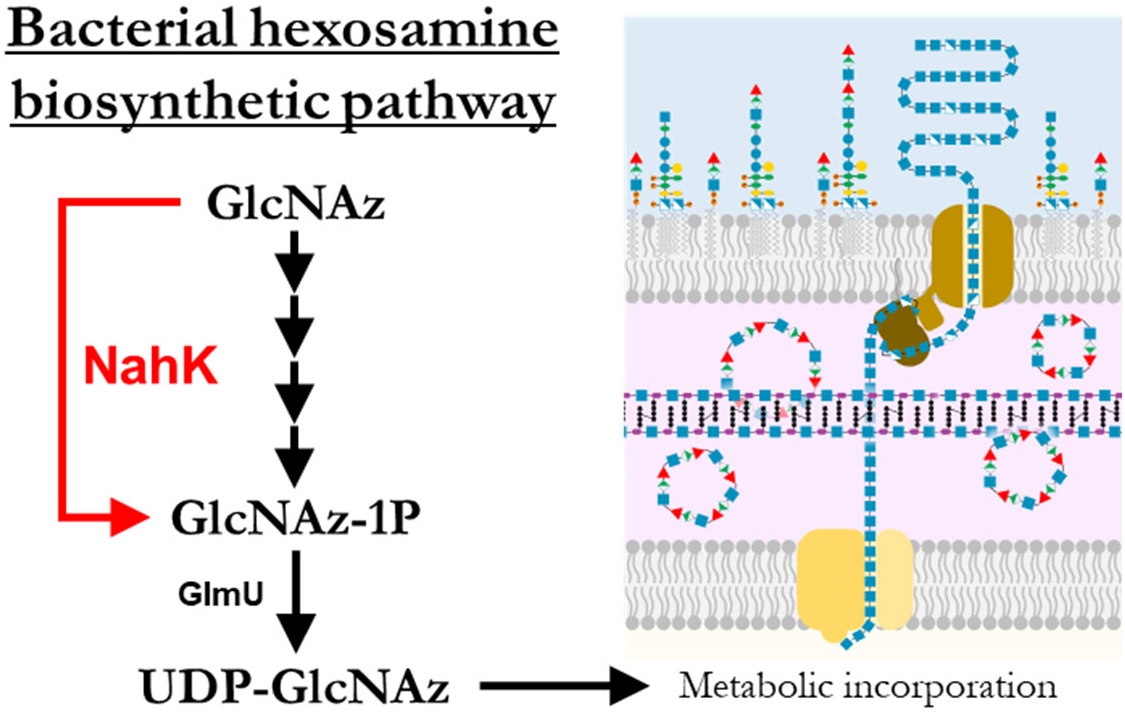

## INTRODUCTION

The envelopes of bacterial cells contain glycans essential for viability, where they maintain cellular integrity, while other glycans are relevant to human health and disease through their roles as virulence factors, and diagnostic markers. Metabolic labelling is a tool for studying bacterial glycobiology in which chemical probes are incorporated into glycan structures.^1^ In general, a functionalized analog of a specific monosaccharide is fed to bacteria which then generates the activated nucleotide sugar via endogenous processing. Recognition of the unnatural metabolite as a glycosyltransferase substrate leads to installation in a target glycan.

In bacteria, there are two general strategies for metabolic glycan labelling. First, using an uncommon monosaccharide, or a monosaccharide in a dedicated glycan synthesis pathway, allows for incorporation into a narrow range of target glycans. For example, the prokaryotic sugar bacillosamine is found in the glycoproteins of certain species, allowing selective labelling of these organisms in a diverse microbial community.^2^ Analogues of rare deoxy amino L-sugars such as *N*-acetylrhamnosamine and *N*-acetylquinovosamine can also be tracked to their glycan destinations to discriminate between bacterial species and/or strains.^3^ Additionally, synthetic precursors for sialic,^4^ pseudaminic,^5,6^ and legionaminic acids^7,8^ have been shown to incorporate into their respective glycoconjugates despite requiring several enzymatic steps to generate the nucleotide sugar donors. For a more detailed discussion on chemical reporters for bacterial glycans, the reader is directed to a recent review.^9^

Often the desired transformations to yield a target nucleotide sugar analogue *in cellulo* are not present. To circumvent this challenge, pathway engineering in *Escherichia coli* has enabled labelling with fucose analogues by replacing the native GDP-fucose pathway with a salvage pathway from *Bacterioides fragilis*.^10^ Metabolic engineering has also been applied to the installation of *N*-acetylmuramic acid (MurNAc) derivatives in peptidoglycan assembly using a recycling pathway from *Pseudomonas putida.*^11^

In contrast to targeting a specific glycan for labelling, it is possible to choose a monosaccharide that is central to many glycans and leverage the importance of the metabolite to obtain information on a range of pathways. This approach requires additional validation due to the number of possible labeled structures that can be formed. Analogues of GlcNAc are an interesting choice as a metabolite to explore multiple glycan pathways, as GlcNAc is an efficient food source^12^ and is also involved in the assembly of peptidoglycan, lipopolysaccharide, exopolysaccharides and capsular polysaccharides. A classic chemical reporter for GlcNAc is the C2-modified *N*-azidoacetylglucosamine probe GlcNAz^13^, first used to characterize the *O*-GlcNAc modification in eukaryotic cells^14^. However, later studies reported that UDP-GlcNAz was also epimerized to UDP-GalNAz and incorporated into O-linked glycans, emphasizing the importance of characterizing the fate of monosaccharides in glycan labelling.^15,16^ Additionally, studies in other eukaryotes have demonstrated variability in the usage of GlcNAz, suggesting that metabolic flux or transferase selectivity’s vary sufficiently to alter the distribution of incorporation.^17,18^ In bacteria, GlcNAz has been shown to be used by *Helicobacter pylori* to glycosylate flagellin,^19^ and in *Staphylococcus aureus* cell surface azides were detectable through azide-clickable fluorophores with GlcNAz feeding.^20^ These studies demonstrate the viability of targeting bacterial glycoconjugates with GlcNAz, but raise questions about into which glycans GlcNAz is incorporated.

To label glycans with GlcNAz, the key metabolite UDP-GlcNAz must be generated intracellularly. It is not possible to feed GlcNAz to *E. coli* as the GlcNAc salvage pathway involves de-*N*-acetylation of the 6-phospho intermediate (GlcNAc-1P) by NagA (*Figure 1a*). One strategy to circumvent this challenge that was successful in lactic acid bacteria is to feed the acetylated 1-phosphate species which presumably is cell permeable and is deacetylated *in cellulo* before entering the HBP pathway.^21^ However, many bacteria, including *E. coli,* are deficient in the non-specific esterase activity necessary to reveal the active probe, limiting the scope of this strategy.^22^ In an alternative approach modified pathways have been introduced into *E. coli*; for example an engineered glycosyltransferase OleD facilitated the formation of unnatural nucleotide sugars from aryl-glycosides.^23^ Using 2-chloro-4-nitrophenyl GlcNAz in *E. coli* expressing OleD, the authors were able to demonstrate the *in situ* production of UDP-GlcNAz and subsequent incorporation of GlcNAz into peptidoglycan.^24^ In another approach we have used the promiscuous *Bifidobacterial* enzyme NahK to generate a set of C6-modified UDP-GlcNAc derivatives *in cellulo* as glycosyltransferase inhibitors.^25^ NahK is a GlcNAc 1-kinase with a high tolerance for synthetic and modified substrates including GlcNAz,^26,27^ and has been used in chemoenzymatic syntheses of various 1-phosphosugars.^28^ It was hypothesized that GlcNAz could be transformed to UDP-GlcNAz in *E. coli* with endogenous expression of recombinant NahK, and subsequently incorporated into the profusion of amino sugar-containing bacterial glycans.

**Figure 1.**
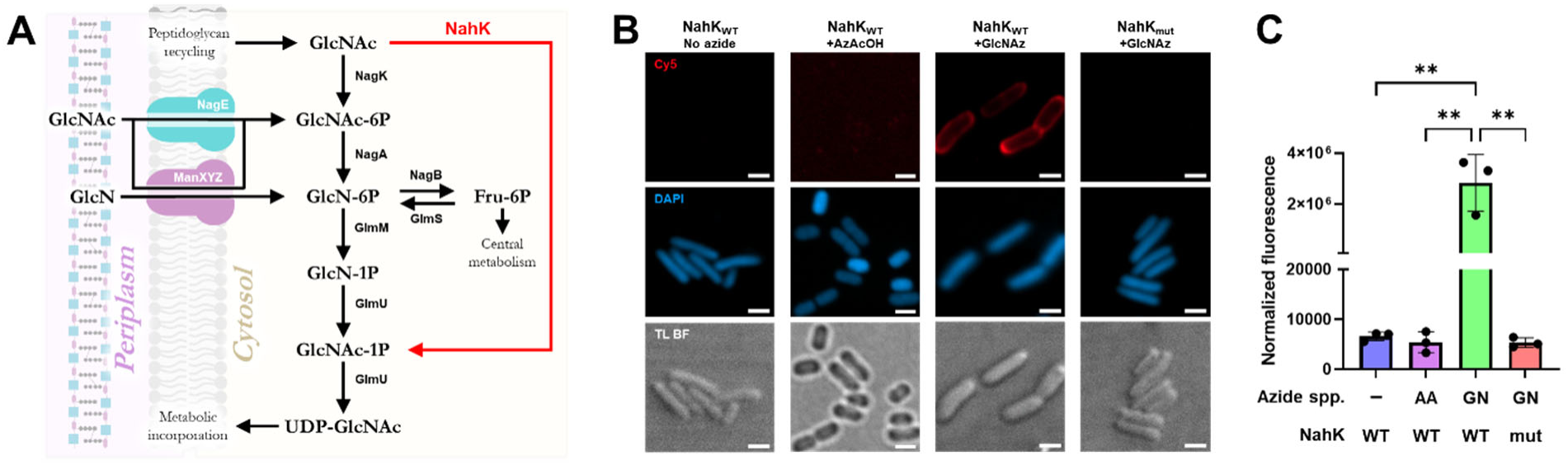
**A**) Hexosamine biosynthetic pathway (in black). GlcNAc and GlcN are imported or recycled from peptidoglycan. The fate of metabolites is decided by the relative activities of NagB and GlmS. Shown in red is the alternative pathway provided by the GlcNAc 1-kinase NahK, allowing GlcNAc to be directly phosphorylated to GlcNAc-1P. **B**) Confocal laser scanning microscopy. *E. coli* BW25113 pBAD(*nahK*) cells were grown (4 hrs at 37 °C) with GlcNAz (1 mM) or AzAcOH (5 mM). Cells were incubated with DBCO-Cy5 (10 µM; 1 hr at 37 °C) and counterstained with DAPI before widefield (TL BF). Experiment was performed twice with similar results. Scale bar is 1 μM. **C**) GlcNAz incorporation into whole cells. *E. coli* BW25113 pBAD(*nahK*) were grown (4 hrs at 37 °C) with azide species: GlcNAz (GN; 1 mM) or AzAcOH (AA; 5 mM). Cells were incubated with DBCO-Cy5 (10 µM; 1 hr at 37 °C) and measured in bulk for retained Cy5 fluorescence after washing; values are normalized to the number of cells per sample. Error bars represent ±SD of 3 biological replicates; significance was determined using unpaired t-tests. **P ≤ 0.002.

Here we sought to define the use of UDP-GlcNAz in *E. coli* K12 based on the well studied glycan biosynthetic machinery which produce the enterobacterial common antigen (ECA), lipopolysaccharide (LPS), and poly-*N*-acetylglucosamine (PNAG). We find the NahK system produces UDP-GlcNAz *in situ*, that GlcNAz is readily incorporated into peptidoglycan, that incorporation into ECA and LPS cannot be detected, and that small amounts are incorporated into PNAG. Discerning the incorporation targets of UDP-GlcNAz in *E. coli* at a molecular level validates the strategy of using sugar 1-kinases to achieve metabolic carbohydrate engineering and as a tool to follow glycan assembly in *E. coli*.

## RESULTS

### *In vivo* incorporation

The incorporation of GlcNAz into *E. coli* BW25113 cells via the alternative hexosamine salvage pathway introduced by the expression of the hexosamine-1-kinase (NahK) from *Bifidobacterium longum* JCM1217 using a pBAD expression system was evaluated^25^. Cells expressed either wild-type NahK (NahK_WT_) or an inactive mutant (NahK_mut_) as a negative control for NahK activity. The incorporation of azides into the bacteria was visualized through strain-promoted alkyne-azide cycloaddition (SPAAC) with dibenzocyclooctyne-Cy5 dye (DBCO-Cy5) by confocal microscopy. Cy5 labelling of *E. coli* is confined mainly to the cell periphery and is both GlcNAz- and NahK-specific; bacteria expressing the inactive NahK_mut_ do not display any pronounced peripheral staining (*Figure 1B*). To confirm that a potential product of GlcNAz catabolism, 2-azidoacetic acid (AzAcOH), was not the source of azide labelling, AzAcOH was added to the cultures, but no Cy5 labelling was observed, suggesting that no significant transfer of 2-azidoacyl groups from GlcNAz to the cell surface is occurring. To complement the microscopy results, bulk cell fluorescence analysis was carried out and gave similar NahK and GlcNAz-dependent labelling (*Figure 1C, Figure S1*). The degree of labelling was dependent on the GlcNAz concentration (*Figure S2*) however at higher concentrations decreases in cell growth were observed (*Figure S3*). Microscopy of labelled cells grown at high GlcNAz (>1 mM) concentrations revealed unusual cell morphology consistent with disrupted peptidoglycan biosynthesis (data not shown).

### Nucleotide sugar detection

The whole cell measurements demonstrated successful incorporation of azide species in a NahK-dependent fashion. It was expected that this was occurring via the hexosamine processing pathway, where the product of NahK, GlcNAz-1-phosphate (GlcNAz-1P), would be converted by GlmU to the central metabolite UDP-GlcNAz for subsequent glycan synthesis (*Figure 1a*). To support this hypothesis, GlcNAz-fed cells were assayed for the presence of UDP-GlcNAz. The initial attempt to identify the unnatural metabolite involved a previously described ion-pair reverse phase HPLC experiment with tetrabutylammonium phosphate as the ion-pairing reagent.^25^ However, UDP-GlcNAz has an unexpectedly long retention time using these conditions (*Figure S4*) leading to peak broadening, and using higher mobile phase concentrations to reduce elution time led to rapid column degradation when running metabolite extracts (data not shown).

Gel electrophoresis was seen as a promising alternative method as it has been applied to the analysis of carbohydrates including nucleotide sugars such as UDP-GlcNAc.^29^ Two fluorophores were investigated for visualization of the GlcNAz derivatives by PAGE in an approach similar to fluorescence assisted carbohydrate electrophoresis (FACE). Labelling with DBCO-Cy3; negatively charged, or DBCO-TAMRA; neutral charge, imparts different electrophoretic mobilities to the GlcNAz derivatives in the PAGE analysis (*Figure S5*). Both fluorophore conjugates readily resolved GlcNAz, GlcNAz-1P and UDP-GlcNAz, however TAMRA labelling gave moderately better results due to less background fluorescence entering the gel and only negatively-charged species migrating (*Figure 2A*). Additional confirmation that visualized bands represent phosphorylated GlcNAz derivatives comes through acid-catalyzed hydrolysis of the sugar-phosphate bond, converting the phosphorylated species to tagged GlcNAz.

**Figure 2.**
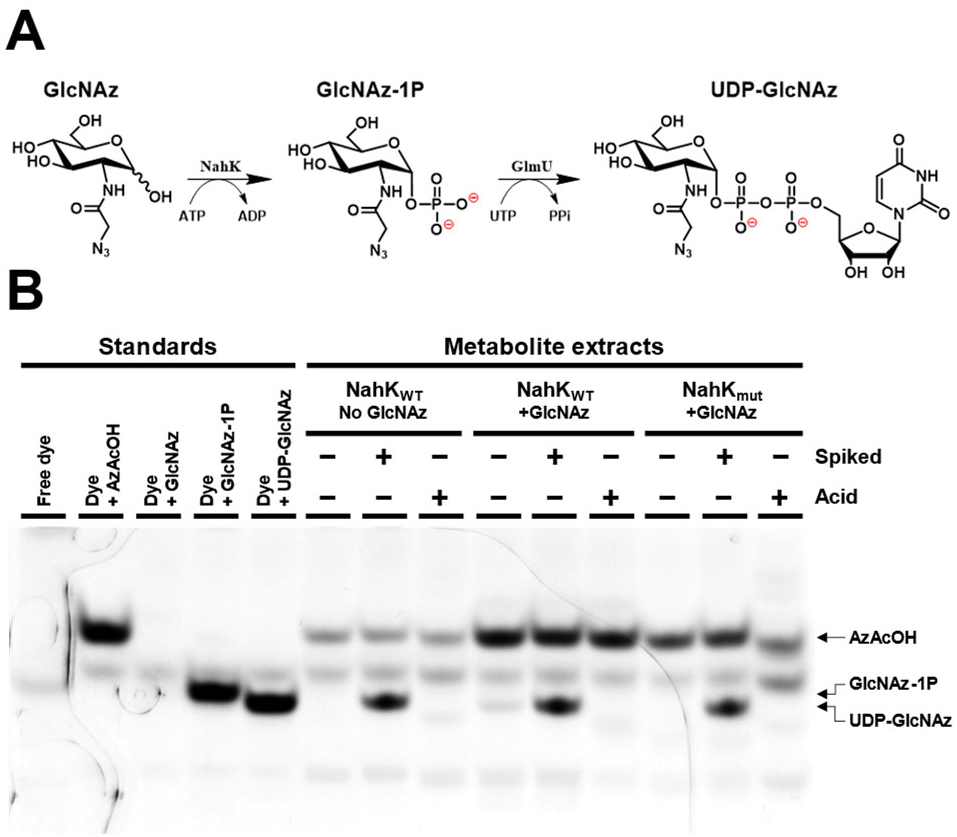
**A**) GlcNAz metabolic processing. GlcNAz imported into the bacterial cytosolic space is phosphorylated by the recombinant NahK kinase to generate GlcNAz-1P, which is then recognized by the uridyltransferase domain of the essential enzyme GlmU leading to *in cellulo* production of UDP-GlcNAz. **B**) FACE gel of metabolite extract. Nucleotide sugar extracts of GlcNAz-fed *E. coli* BW25113 pBAD(*nahK*) cells (2-3 hrs at 37 °C; 1 mM GlcNAz) were selectively heated with acid to hydrolyze sugar phosphates (**Acid**), incubated with DBCO-TAMRA to tag azides (10 µM; 16 hrs at 37 °C), and spiked with UDP-GlcNAz migration standard (**Spiked**). FACE gel was run with migration standards of tagged azide species and imaged for TAMRA fluorescence. Note, TAMRA is neutral at gel running pH and only negatively charged species migrate. Arrows indicate migration distances of various tagged azide species.

Crude metabolite extracts of mid-log phase *E. coli* cells were collected and selectively treated with acid before incubation with DBCO-Cy3 or TAMRA. Lanes with extracts obtained from GlcNAz-fed cells show a faint band with similar migration to the UDP-GlcNAz standard lane (*Figure 2B*, *Figure S6*). This band overlaps a sample spiked with UDP-GlcNAz and is absent in cell extracts that lack GlcNAc-1-kinase activity (NahK_mut_ lanes). In samples pretreated with acid to hydrolyze glycosidic phosphate bonds the UDP-GlcNAz band is lost, with the accompanying emergence of a tagged-GlcNAz band seen in FACE gels using Cy3 labelling (*Figure S*6). Other unidentified signals are also present in these analyses. Most notably, a band corresponding to a species with a similar migration to tagged AzAcOH appears in all cell extract samples, even those not exposed to any azide containing species. This band likely represents multiple species and may contain AzAcOH or other azide derivatives such as GlcNAz-6P, as well as non-azide species that react with DBCO. In an effort to reduce the complexity of gel lanes, methoxypolyethylene glycol azide (PEG-Az) was added to gel samples following tagging with dye, to sequester any unreacted dye or other azide-reactive components of the sample and retard their migration through the gel (*Figure S7*). No major deconvolution was achieved through this strategy, demonstrating the excess dye is likely reacting through non-specific pathways with other sample components and generating unwanted side products^30^.

### Peptidoglycan incorporation

Given the cytosolic UDP-GlcNAz generation, and the clear azide-dependent labelling in *E. coli,* the key question was into which glycans are the azides incorporated? A major destination for GlcNAc in bacteria is the peptidoglycan; both monosaccharides in the alternating GlcNAc(β1→4)-*N*-acetylmuramic acid (MurNAc) backbone are derived from GlcNAc (*Figure S8*). This repeating unit is generated as a lipid-linked intermediate (Lipid II) on the cytosolic leaflet of the cell membrane before being flipped and transferred to the nascent peptidoglycan chain.^31^ As has been previously shown in OleD-expressing *E. coli*, UDP-GlcNAz is used by MurG to generate Lipid II.^24^

To confirm the presence of GlcNAz in PG through the NahK system, sacculi were isolated from GlcNAz-treated bacteria possessing NahK activity at mid-late log phase.^32^ Peptidoglycan samples, both as whole sacculi and lysozyme treated samples, were labelled with DBCO-Cy5 and visualized on polyacrylamide FACE gels (*Figure 3*, *Figure S9*).

**Figure 3.**
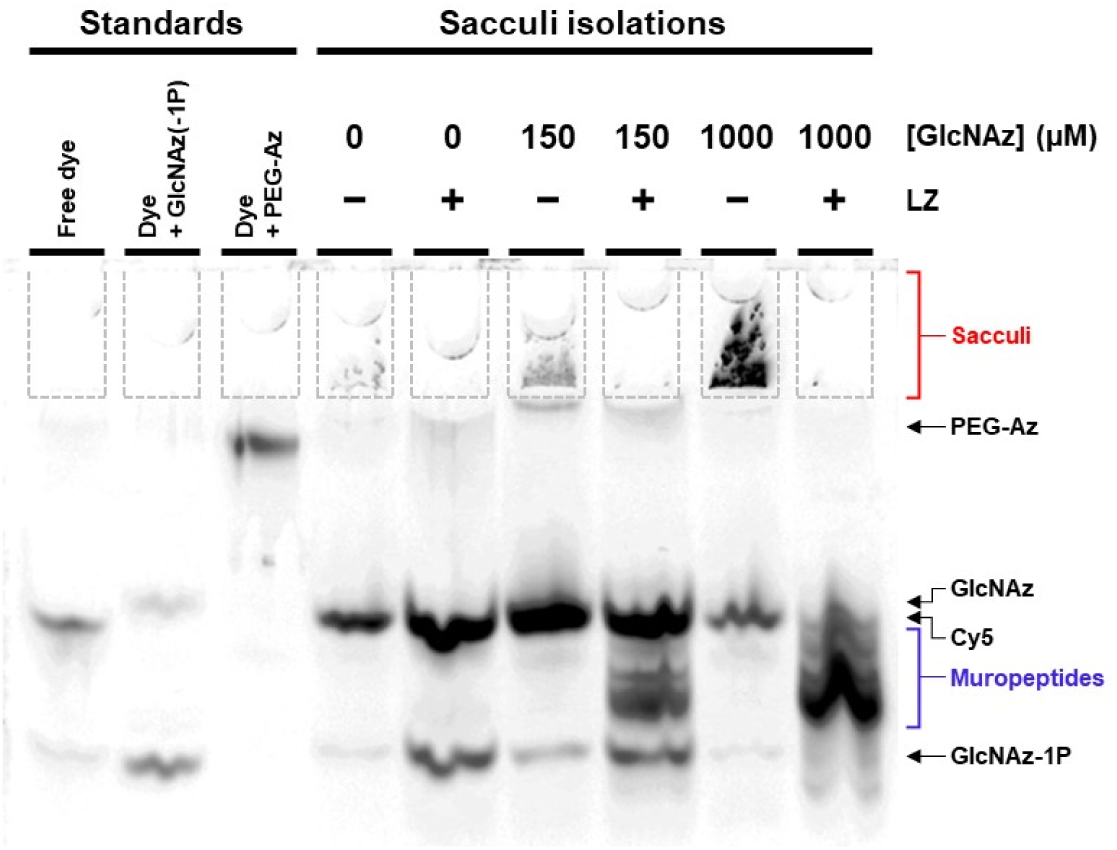
FACE gel of peptidoglycan sacculi. Isolated sacculi (marked in red) from GlcNAz-fed *E. coli* BW25113 pBAD(*nahK*) cells (6 hrs at 37 °C) were selectively treated with lysozyme (**LZ**; 40 mg/mL; 16 hrs at 37 °C) to generate muropeptides (marked in blue) and incubated with DBCO-Cy5 to tag azides (10 μM; 1 hr at 37 °C). FACE gel was run with migration standards of various tagged azide species (GlcNAz(−1P) standard was generated as an incomplete NahK reaction with GlcNAz) and imaged for Cy5 fluorescence. Arrows indicate migration distances of various tagged azide species. Sample wells are outlined in dashed lines for ease of viewing.

In non-hydrolyzed samples, the large insoluble sacculi do not migrate into the gel and remain in the sample wells, where they can be seen as a fluorescent mass in samples derived from GlcNAz-fed cells. Lysozyme specifically cleaves the PGN glycan backbone, producing a series of smaller soluble fragments, which appear as a ladderlike pattern of bands running above the standard GlcNAz-1P. The soluble samples were analyzed by ESI-MS but no peaks corresponding to azide modifications were observed, likely indicating low levels of GlcNAz incorporation, however the identity of the PG fragments could be clearly observed.

### Wzx/Wzy-dependent glycans

In the chosen model Gram-negative system, both the enterobacterial common antigen (ECA) and lipopolysaccharide (LPS) can be found. In the trisaccharide repeating unit of ECA, 2 of the 3 sugars originate from UDP-GlcNAc; *N*-acetylmannosaminuronic acid (ManNAcA) and GlcNAc (*Figure S8*).^33^ LPS is built up from a phosphoglycolipid core, lipid A, composed of a GlcN disaccharide pair which is extensively modified with acyl chains (*Figure S8*). Additionally, the outer core in K12 strains is non-stoichiometrically modified with a terminal GlcNAc^34,35^; *E. coli* K12 has no elaborate O-antigen due to the loss of a rhamnosyltransferase and therefore possesses rough-type LPS.^36^ The transfer of GlcNAc to ECA and to the terminal sugar in the LPS outer core proceeds via undecaprenyl-pyrophosphoryl-GlcNAc (also known as bactoprenol-pyrophosphoryl-GlcNAc, GlcNAc-BPP) formed by WecA (also known as Rfe). GlcNAc-BPP initiates the biosynthesis of the repeating units of both ECA and the *E. coli* K12 O-antigen (*Figure 4a*). The lipid-linked subunits are further modified by other glycosyltransferases and flipped from the cytoplasmic to the periplasmic leaflet of the inner membrane, where they are polymerized by the Wzx/Wzy machinery and transferred to the final carrier lipids: core oligosaccharide (OS) to make lipooligosaccharide (LOS) or OS-linked ECA (ECA_LPS_), with ECA repeats also used to generate cyclical glycans (ECA_CYC_) or installed on diacylglycerol (ECA_PG_) through a phosphodiester linkage.^33^

**Figure 4.**
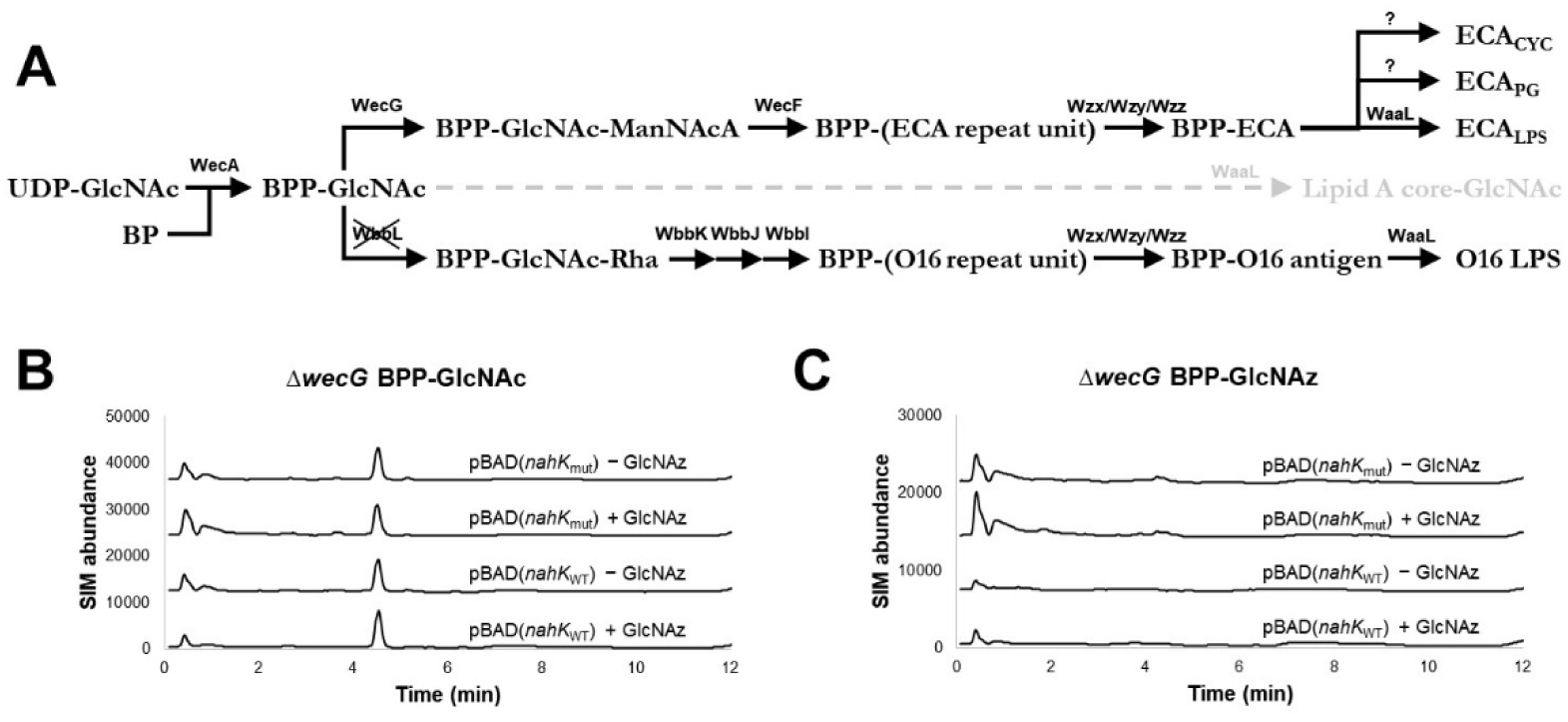
**A**) Biosynthesis of Wzx/Wzy-dependent glycans. The transferase WecA generates the shared lipid-linked GlcNAc precursor in the synthesis of both ECA and O-antigen using bactoprenol (**BP**). In *E. coli* K12 strains the rhamnosyltransferase WbbL is not expressed leading to truncation of O-antigen production, but the ligase WaaL is capable of transferring a single GlcNAc onto the Lipid A-core acceptor (in grey). **B**, **C**) RP-LC-MS SIM of BPP-linked intermediates. Lysates from ECA knockout strains of *E. coli* MG1655 pBAD(*nahK*) grown with GlcNAz were analyzed by LC-MS, monitoring (**B**) BPP-GlcNAc and (**C**) BPP-GlcNAz.

The lipid-linked structures of LPS and certain forms of ECA allow a shared isolation and characterization to be conducted.^37^ To investigate GlcNAz incorporation into either structure, *E. coli* BW25113 cells were cultured with GlcNAz, harvested and dried *in vacuo* in preparation for phenol-chloroform-pentanes (PCP) extraction of ECA and rough-type LPS (*Figure S10*).^38^ An additional K12 strain, AB113 *rfe*::Tn10 is deficient in ECA production and terminal GlcNAc installation onto core OS, and was used as a negative control for glycan biosynthesis.^39^ The resulting extracts were labelled with DBCO-Cy5 and analyzed via SDS-PAGE (*Figure S11*). Comparing the fluorescence imaging of the gels and the silver-stained signals, there is no evidence of any DBCO-Cy5 labelling, suggesting that UDP-GlcNAz is not tolerated as a substrate for glycan incorporation, or that GlcNAz-containing glycans are exceedingly low abundance.

To determine which step in the pathway failed to use the GlcNAz derivative, the BPP conjugates were analyzed in a Δ*wecG* strain which is known to accumulated GlcNAc-BPP.^40^ Overnight cultures were lysed and analyzed by reverse phase high performance liquid chromatography−mass spectrometry (RP-LC-MS) in selective ion monitoring (SIM) mode for either GlcNAc-BPP (m/z=1128.7) or GlcNAz-BPP (m/z=1156.8) (*Figure 4.*). Additional controls display unmodified bactoprenol (BP) acceptor (m/z=845.7) (*Figure S12*). These experiments clearly showed the expected presence of GlcNAc-BPP but failed to show ions consistent with GlcNAz-BPP in the presence of NahK and GlcNAz, agreeing with WecA failing to generate GlcNAz-BPP in this system.

### Biofilm exopolysaccharide

Another likely site of GlcNAz incorporation is the exopolysaccharide poly-β-1,6-*N*-acetylglucosamine (PNAG). This polysaccharide is produced by a variety of distantly related microorganisms,^41^ and is best known for its role as a matrix polymer in bacterial biofilms.^42^ PNAG is partially deacetylated (*Figure S8*), giving it an overall positive charge which is thought to aid in its role as a biofilm matrix component by promoting cell-to-cell and cell-to-surface contacts.^43^

In *E. coli* and other Gram-negative organisms PNAG is produced on the cytosolic side of the cell membrane by the glycosyltransferase complex PgaCD from UDP-GlcNAc which polymerizes and translocates the growing chain across the membrane.^44^ A related enzyme complex exists in Gram-positive organisms.^45^ The nascent PNAG chain is then partially-deacetylated before being exported to the extracellular environment to form the biofilm matrix. C6-modified UDP-GlcNAc analogues are incorporated into PNAG and lead to truncation of the polymer, however the specificity of the synthases for the acetamido moiety has not been evaluated.^25^

To ascertain whether GlcNAz could be incorporated into cell-produced PNAG, a whole cell approach with a Δ*csrA* PNAG-overproducing strain of *E. coli* was used.^46^ GlcNAz and NahK activity did not appear to affect the development of static biofilms as analyzed by crystal violet biomass staining (*Figure S13*). After growth of *E. coli* MG1655 *csrA::kanB* strains in liquid culture, PNAG was extracted using treatment with concentrated ammonium bicarbonate which was found to readily solubilize the PNAG. Previously high salt concentrations have been found to solubilize PNAG, and we found the volatile salt worked similarly^47^ (*Figure S14*). Successful PNAG isolation was confirmed through a dot blot assay using a lectin-peroxidase conjugate (WGA-HRP) with limited signal observed from strains that do not overproduce PNAG or upon treatment with PNAG-specific hydrolase Dispersin B (DspB).^25^ Extract from an *E. coli* strain that does not overproduce exopolysaccharide fails to generate blot-based luminescence, while the positive signal of other extracts is mostly eradicated upon treatment with the PNAG-specific hydrolase Dispersin B (DspB)^48^; this is not the case upon treatment with the non-specific protease Proteinase K (ProtK) (*Figure S15*). Analyzing the extracted material via SDS-PAGE showed protein contamination, including abundant membrane protein OmpC (*Figure S16*),^49^ necessitating a protease treatment to deconvolute PNAG staining on polyacrylamide gels.^47^

Stationary cell cultures of *E. coli* MG1655 *csrA::kanB* grown with GlcNAz were washed, the PNAG was extracted and analyzed via dot blot (*Figure 5a*). Upon seeing no significant difference in PNAG extracts between Δ*csrA* cell cultures in the presence of GlcNAz, samples were proteolyzed, selectively treated with DspB, and labelled with DBCO-Cy5 before PAGE analysis (*Figure 5b,c*). The amount of PNAG material stained via Coomassie Brilliant Blue is seen to vary extraction-to-extraction however bulk quantification of PNAG amounts between strains was not seen to vary. The sole sample displaying fluorescent signal is the extract derived from cell cultures fed with GlcNAz and expressing active NahK, and hydrolysis with DspB appears to increase the electrophoretic mobility of this signal, suggesting the smaller molecular weight oligosaccharides are being generated.

**Figure 5.**
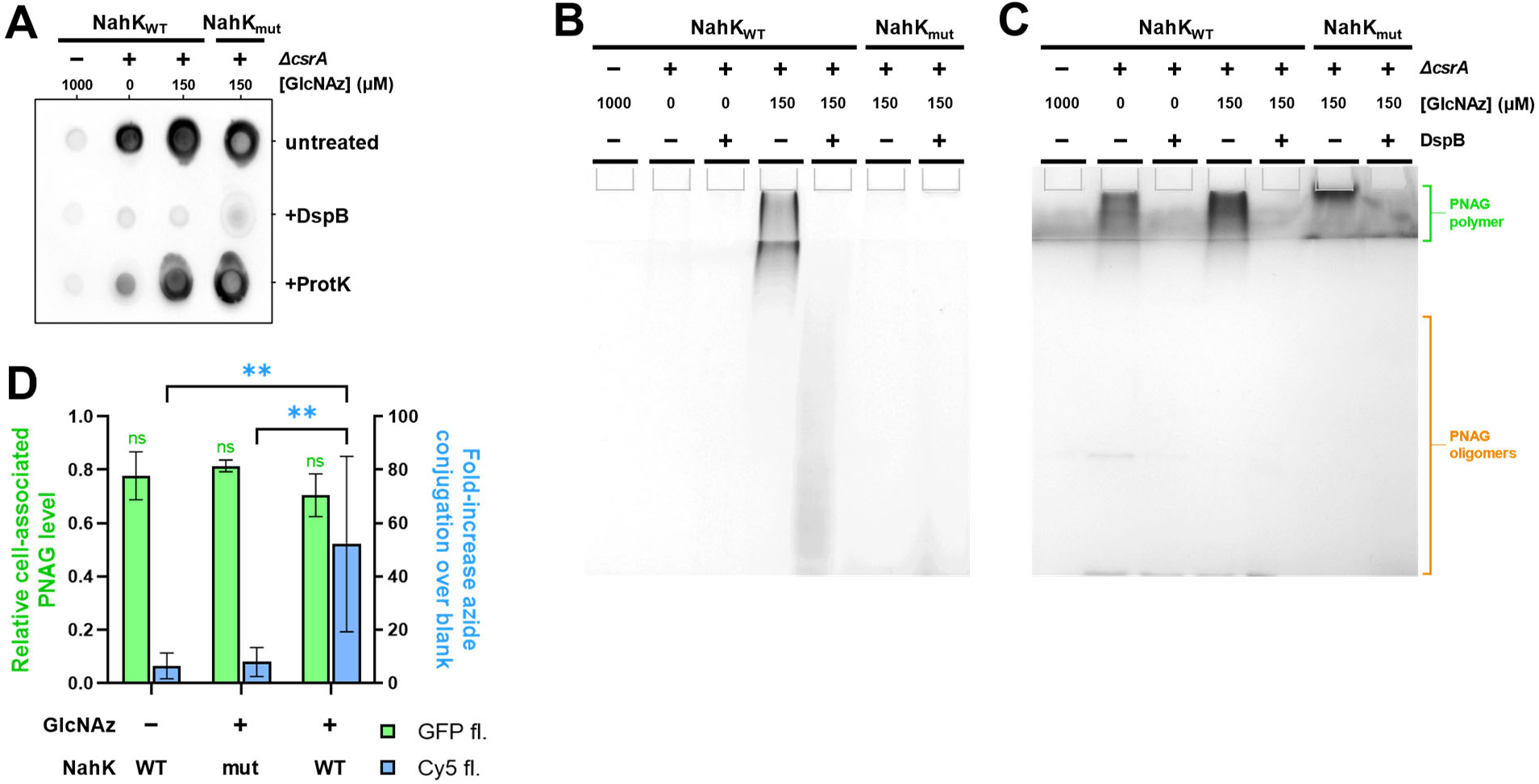
**A**) PNAG extract dot blot. PNAG extracts from *E. coli* pBAD(*nahK*) cultures (2 days at rt) were spotted on nitrocellulose membranes and probed with WGA-HRP. Select samples were pretreated with the PNAG-specific hydrolase DspB (5 μM; 4 hrs at 37 °C), or the broad-spectrum protease ProtK (1 mg/mL; 4 hrs at 37 °C). Overproducing strains of PNAG (Δ*csrA*) generate more signal than the wildtype control strain. **B**, **C**) PNAG extract PAGE. Deproteinized PNAG extracts from GlcNAz-fed *E. coli* pBAD(*nahK*) cultures (2 days at rt) were labelled with azide-reactive DBCO-Cy5 (10 μM; 1 hr at rt) and analyzed by gel electrophoresis using (**B**) Cy5 fluorescence and (**C**) Coomassie stain. Select samples were pretreated with the PNAG-specific hydrolase DspB; effect is apparent as a disappearance of the slow-migrating species (marked in green) and generation of a higher-migration species (marked in orange). **D**) Cell-associated PNAG assay. GlcNAz-fed *E. coli csrA*::*kanB* pBAD(*nahK*) cells (4 hrs at 37 °C) were labelled with DBCO-Cy5 to tag azides (10 μM; 1 hr at rt), and incubated with PNAG-binder GFP-DspB^E184Q^. Cells were washed with buffer and DspB enzyme to hydrolyze PNAG. Data plotted represents the GFP and Cy5 signals in the DspB-solubilized buffer fraction. GFP values are normalized to the fluorescence of the initial labelling solution; Cy5 values are relative to a blank sample without added dye. Error bars represent ±SD of 3 biological replicates; significance was determined using unpaired t-tests. **P ≤ 0.0037, ns = not significant.

As further evidence for the incorporation of GlcNAz into PNAG, a previously-developed assay for quantifying cell-surface PNAG levels was adapted to measure GlcNAz incorporation levels (*Figure 5d*).^50^ Briefly, cells at mid-log were harvested and labelled with both DBCO-Cy5 and a GFP-tagged PNAG-binding probe. Sequential washes in 0.22 µm spin filter microcentrifuge tubes then allow for enzymatic treatments with total retention of cells and analysis of the resulting solution. Treatment with DspB will hydrolyze cell-surface PNAG, resulting in the release of both Cy5 and GFP fluorescence into the filtrate; the former an indicator of GlcNAz incorporation, the latter a measure of total PNAG amount. All samples displayed similar levels of PNAG production, but the cells expressing active NahK enzyme also showed high Cy5 fluorescent signal when fed GlcNAz during culturing (*Figure S17*).

Following detection of fluorophore-conjugated PNAG species, attempts were made to obtain mass spectral support for incorporation. However in the DspB treated samples only unmodified PNAG oligomers were observed (*Figure S18*).

### *In vitro* PNAG incorporation

To shed light on low incorporation levels of GlcNAz into PNAG, an *in vitro* PNAG biosynthesis system with the *E. coli* synthase complex was explored. Using a previously described membrane preparation^25^ of the *E. coli* PgaCD synthase complex,^51^ UDP-GlcNAz was tested as a substrate for *in vitro* polymerization. The unnatural nucleotide sugar was chemoenzymatically synthesized from GlcNAz following a previously reported protocol,^28^ and PNAG product formation in reaction mixture aliquots was visualized using dot blots (*Figure 6a*).

**Figure 6.**
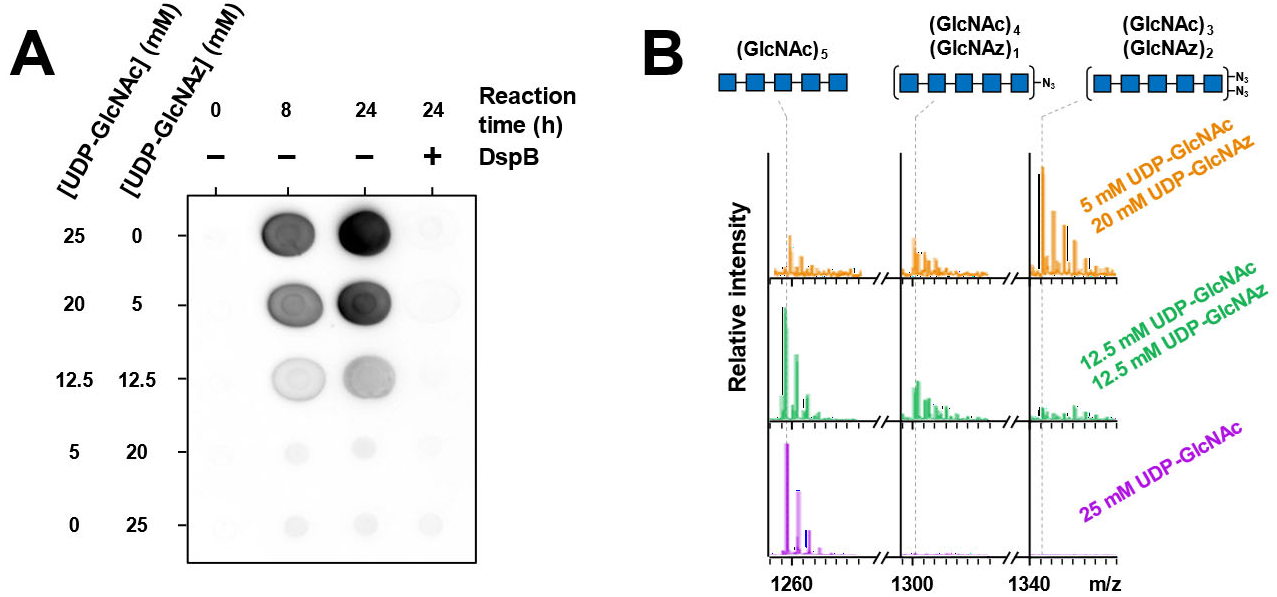
**A**) PgaCD polymerase reaction. Using varying ratios of UDP-GlcNAc and -GlcNAz as sugar donors, reaction mixtures incubated with PgaCD membrane preparations (24 hrs at 37 °C) were spotted on nitrocellulose membranes. Reaction endpoints were also selectively treated with DspB to hydrolyze PNAG (5 µM; 4 hrs at 37 °C), thereby resulting in an apparent loss of product. WGA-HRP was used for visualization of reaction progress. **B**) ESI-MS of PgaCD reaction. PgaCD reaction mixtures were digested with DspB (5 µM; 90 min at 37 °C). Sodiated pentasaccharide ions [M+Na]^+^ containing 0, 1, or 2 GlcNAz units are displayed (see SI for full spectra of each reaction). Peak intensities are relative to the highest intensity signal in each respective spectrum.

The signal from a 24-hour reaction is completely eradicated upon treatment with the DspB hydrolase, confirming the PNAG origin of the luminescent signal. Extending the reaction to further timepoints did not give an increase in product (data not shown). When present as the sole sugar nucleotide, UDP-GlcNAz does not appear to be a competent substrate. Samples with combinations of UDP-GlcNAc and UDP-GlcNAz show product formation, however UDP-GlcNAz appears to inhibit product formation. The binding selectivity of WGA for GlcNAz is not known but it is expected that the lectin would continue to give a faithful measurement of PNAG containing some GlcNAz, as WGA is capable of recognizing single units of terminal or internal GlcNAc in a polymer.^52^ The inhibitory behaviour of UDP-GlcNAz indicates it is a poor substrate and potentially weak inhibitor of PgaCD (*Figure S19*).

To detect the potential low levels of incorporation of GlcNAz into PNAG, reaction mixtures were partially hydrolyzed with DspB to obtain soluble oligosaccharides prior to fractionation on a graphitized carbon column. Digested samples were submitted directly for ESI-MS analysis. Mass spectral signals consistent with oligosaccharides containing GlcNAz were observed confirming that UDP-GlcNAz is a poor but tolerated substrate of PgaCD (*Figure 6b*, *Figure S20*). As the UDP-GlcNAz:UDP-GlcNAc ratio of the reaction was increased, the number of GlcNAz units present per oligosaccharide is also seen to increase in a concentration-dependent manner, however a clear preference for UDP-GlcNAc can be observed by the limited GlcNAz incorporation at equivalent donor concentrations. (*Figure S21*, *Figure S22*, *Figure S23*, *Figure S24*, *Figure S25*).

## DISCUSSION

### GlcNAz in the hexosamine pathway

The route of GlcNAz uptake to the cytosol in *E. coli* is a subject of ongoing studies, but the amount available for conversion to GlcNAz-1P by NahK must be sufficient to allow for competition with endogenous GlcNAc-1P production even at lower feeding concentrations (∼250 μM). Cytosolic GlcNAz could be undergoing processing by the GlcNAc-6-kinase NagK and deacetylase NagA via the native HBP pathway, leading to production of GlcN-1P and AzAcOH, however AzAcOH is unlikely to be a major contributor to the degree of cell-envelope azide incorporation, as control experiments with AzAcOH did not lead to azide labelling. Additionally, the structure of NagA shows a constrained binding pocket for the acetamido group, making it unlikely that GlcNAz-6P would be accepted as a substrate.^53^ Conversely the co-crystal structure of GlcNAc bound to a NagK ortholog does appear to permit bulkier substituents at the C2-position suggesting GlcNAz-6P maybe produced.^54^

### GlcNAz in bacterial labelling

A series of strategies have been developed to label the peptidoglycan of bacteria and the NahK/GlcNAz combination described here offers a useful complementary method. MurNAc can be conveniently labeled in PG using MurNAc derivatives^11^ and unnatural D-amino acids^55^ can be used to label the peptide side chains. Here we show that GlcNAz derivatives have a strong preference for labelling the peptidoglycan over other GlcNAc containing glycans in *E.coli* K-12. The use of NahK expressing cells provides an alternative means to achieve sufficient UDP-GlcNAz concentrations in the cytosol to the previously reported use of glycosyl transferase OleD in combination with a phenolic GlcNAz glycoside.^24^ Unlike the feeding strategy using the free sugar, the production of UDP-GlcNAz from the glycoside also liberates the corresponding nitrophenol during OleD reaction, potentially leading to buildup of a toxic compound.^56,57^ Additionally, the turnover of GlcNAz-labelled peptidoglycan likely differs between OleD or NahK-expressing bacteria, as the peptidoglycan fragments released by the lytic transglycosylases are imported and metabolized to GlcNAc.^58^ Thus, GlcNAz could be recycled to UDP-GlcNAz in the presence of NahK, and be reincorporated. In contrast, no pathway is available in the OleD system to recycle cytosolic GlcNAz.

UDP-GlcNAz was also incorporated into PNAG by *E. coli* albeit at low levels due to the preference of PgaCD for GlcNAc over GlcNAz. Interestingly, *in vitro* reactions without added UDP-GlcNAc with UDP-GlcNAz as the soul substrate for PgaCD, showed mass spectra corresponding to oligosaccharides with a mix of both GlcNAc and GlcNAz. This may be due to residual UDP-GlcNAc or PNAG oligomers carried over in the enzyme purification. The latter explanation is consistent with slow release of bound-PNAG oligomers, as has been previously suggested in inhibition studies using chain terminating inhibitors.^25^ Furthermore, there were no oligomers observed that contain exclusively GlcNAz suggesting UDP-GlcNAz is unable to initiate the polymer synthesis. The *in vitro* experiments with PgaCD using defined nucleotide sugar concentrations give informative bounds on the likely cytosolic ratio of UDP-GlcNAc to UDP-GlcNAz. In vitro, it was challenging to detect GlcNAz incorporated PNAG by ESI-MS at donor ratios below 1:1, and no incorporation was detected *in vivo* suggesting the concentration of UDP-GlcNAz must be significantly below that of UDP-GlcNAc. In practical terms the low level of UDP-GlcNAz incorporation into PNAG, and the small amounts of PNAG produced by *E. coli* in the absence of biofilm formation suggest that for most applications this incorporation will not be significant.

The substrate specificity of WecA has not previously been investigated despite the central role this enzyme plays in the synthesis of Wzx/Wzy-dependent glycans ECA and O-antigen in many bacterial strains. The lack of recognition of GlcNAz suggests WecA has a narrow substrate scope. It is tempting to speculate that the specificity may be due to efforts by the bacteria to avoid toxicity that may occur if unusable lipid linked intermediates accumulate diminishing the pool of undecaprenol.^59^ It is important to note that in other bacterial strains GlcNAc is introduced into the O-antigen repeat at positions other than the reducing terminus, for example *E.coli O3* O-antigen contains GlcNAc within the repeat.^60^ The substrate specificity of the glycosyltransferases introducing these GlcNAc residues will need to be determined before one can rule out incorporation of GlcNAz into O-antigen in these strains.

### Other GlcNAc modifications

*E. coli* uses UDP-GlcNAc as the starting metabolite to generate UDP-ManNAcA. The epimerase WecB first catalyzes the conversion to UDP-ManNAc, which is used by the dehydrogenase WecC to make UDP-ManNAcA. This sugar nucleotide is then used in the biosynthesis of ECA. This would provide a route for UDP-GlcNAz to be incorporated in ECA as ManNAzA, avoiding the need for WecA transferase activity. Previous literature however suggest that the unnatural structure would again run into substrate tolerance issues, as a homologue of WecB in *Neisseria meningitidis* was found to be unable to epimerize the C2 position of UDP-GlcNAz.^61^ Other monosaccharides derived from GlcNAc are possible labelling targets, for example GalNAz potentially derived by an epimerase could lead to incorporation into O-antigens containing GalNAc such as *E. coli* O23A.^60^ With the knowledge that WecA does not accept GlcNAz it would be possible to interrogate the incorporation of GlcNAz into these strains.

## CONCLUSION

Through engineering an alternate metabolic route in the bacterial hexosamine biosynthetic pathway, the utilization of the well-known GlcNAc metabolic probe GlcNAz has been characterized in an *E. coli* K12 strain, the most widely used laboratory strain of bacteria. In addition to validating the tolerance of peptidoglycan assembly machinery towards GlcNAz as a substrate, the PNAG biosynthetic complex was also found to be capable of using the unnatural sugar for labelling PNAG. A new gentle extraction method for PNAG was also described. Furthermore, the intolerance of WecA towards GlcNAz demonstrates that modification at the C2 position is not tolerated, informing future development of carbohydrate-based metabolic probes or antimicrobial compounds destined for ECA or O-antigen. The ability to generate unnatural UDP-GlcNAz *in cellulo* starting from the free sugar could be used in other organisms, with the same well-conserved HBP, or used to investigate other amino sugar-containing glycans. Furthermore, this strategy could be used to develop functional polysaccharides for use in biotechnology applications.

## SUPPORTING INFORMATION

Additional experimental details, materials, and methods are included in the SI.

## ACKNOWLEDGEMENTS

Many thanks to the lab of Dr. Chris Whitfield for the gift of an *E. coli rfe::*Tn10 strain, Matthew Jorgenson University of Arkansas Medical Sciences for the ECA knockout strains of *E. coli* and to Dr. Matt Forbes and Chung Woo Fung of the Advanced Instrumentation for Molecular Structure (AIMS) mass spectrometry facility for helpful advice regarding sample analysis.

## AUTHOR CONTRIBUTIONS

A.E., Z.A.M., J.M.T and M.N.: conceptualization. A.E., M. K. D. and A.S.S.: investigation. A.E., and M.N.: writing─original draft. A.E., M.N. J. M T., P.L.H: writing─review and editing. M.N. J. M. T and P.L.H.: supervision. M.N., J.M.T and P.L.H.: funding acquisition. M.N.: project administration.

## Supporting information

### METHODS

Unless otherwise mentioned, all materials and reagents were purchased from Sigma-Aldrich, Bio-Rad, or BioShop Canada. Plate reader measurements were performed using a CLARIOstar instrument from BMG Labtech. Gel and blot imaging was performed using a G:BOX Chemi XT4 (Syngene) imager. ESI-QTOF mass spectrometry data was acquired using an Agilent 6538 UHD mass spectrometer by Dr. Matthew Forbes and Chung Woo Fung at the AIMS facility, Department of Chemistry, University of Toronto.

#### General bacterial culturing methods

All bacterial starter cultures were made up with LB(Miller) media containing appropriate antibiotics, and were inoculated from single colonies grown on LB-agar. Starter cultures were typically incubated overnight at 37 °C with shaking (180 rpm), resulting in saturated cultures. Prewarmed LB media was inoculated with overnight starters at a dilution of 1:100 inoculum/media.

#### Fluorescence-assisted carbohydrate electrophoresis (FACE)

High percentage polyacrylamide gels were prepared as one continuous concentration of acrylamide (35%) with a Tris buffer system (375 mM Tris pH 8.8, 87.5 mM NaCl). It was found that 1.5 mm thick gels gave the most reproducible results, although care must be taken to ensure they do not overheat during casting through the use of precooled solutions. Samples for FACE analysis were typically incubated with azide-reactive (SpAAC) fluorescent dye (10 µM; >1 hour at 37 °C) and mixed 1:1 with 2× FACE loading buffer (375 mM Tris pH 8.8, 40% (v/v) glycerol); 5 μL gel samples were loaded into sample wells. For PEG-labelling, samples already labelled with fluorophore were incubated with PEG-Az (Sigma 689475; 0.5 mM) overnight at 37 °C before being mixed with loading buffer. Gels were electrophoresed at a constant voltage (300 V; 3 hours) in tank running buffer lacking both detergent and reducing agent. The heat generated by the high voltage required to force migration of small analytes is sufficient to begin melting the gel matrix when the apparatus was left at room temperature, so all runs were conducted with the gel box sat in an ice-water bath placed in a fridge (4 °C). Fluorescence images of gels were acquired before removal of the gels from between their glass plates, using a G:BOX Chemi XT4 (Syngene) imager.

#### Synthesis of azidosugars

GlcNAz was synthesized in gram-scale quantities as previously described. GlcNAz-1P and UDP-GlcNAz were generated enzymatically using purified recombinant NahK and GlmU respectively.^1^

#### Purification of PgaCD membrane preparations

Membranes containing recombinantly expressed PgaCD were isolated as described previously.^2^

#### Generation of NahK constructs

The construction of pBAD(*nahK*) expression plasmids has been described previously.^2^

#### Live cell confocal microscopy

LB media (10 mL) with ampicillin (0-100 mg/L) and arabinose (20 mg/L) was inoculated with starter cultures of *E. coli* BW25113 pBAD(*nahK*). Cultures were raised in 50 mL vented culture tubes (180 rpm; 37 °C) for 30 minutes, then GlcNAz (1 mM) or AzAcOH (5 mM) was added to the culture media and the incubation was continued. After ∼3 hours of growth (OD_600_ = 0.5-1.0) an aliquot of cell culture was taken (1 mL) Cell pellets were washed with PBS and resuspended in PBS (0.1 mL). DBCO-Cy5 (10 μM) was added and the samples were incubated in the dark (1 hr at rt). Cells were washed twice more with PBS and resuspended in a DAPI labelling solution (10 µg/mL; 1 mL) before incubation in the dark (10 min at rt). Cells were washed once more with PBS and resuspended (1 mL). Several microlitres of labelled cell suspension was added onto a PBS-based 1% (w/v) agarose pad. When all excess liquid had soaked into the agarose the pad was sandwiched between two sterile acid-washed borosilicate coverslips and the edges sealed with Vaseline-lanolin-paraffin mixture (VALAP, 1:1:1 w/w/w) to prevent evaporation. Microscopy images were obtained with an AxioObserver.Z1 microscope equipped with a Plan-Apochromat 100×/1.40 Oil DIC M27 objective.

#### Whole cell GlcNAz incorporation assay

LB media (1-2 mL) with ampicillin (0-100 mg/L) and arabinose (20 mg/L) was inoculated with starter cultures of *E. coli* BW25113 pBAD(*nahK*). Cultures were raised in 5 mL vented culture tubes (180 rpm; 37 °C) for 30 minutes, then GlcNAz (1 mM) or AzAcOH (5 mM) was added to the culture media and the incubation was continued. Based on OD_600_ measurements from growth curves, 4 hours (OD_600_ = 0.5-1.0) was selected as the optimal incubation timepoint for quantifiable and reproducible levels of incorporation. OD_600_ of culture aliquots (200 μL) were measured in a 96-well microplate, and cells were harvested by centrifugation (21 k rcf; 5 minutes). Cell pellets were washed with PBS and resuspended in PBS to a volume equal to 1/10^th^ the initial culture volume. Azide-reactive DBCO-Cy5 (10 μM) was added and the samples were incubated in the dark (rt; 1 hour). Cells were washed twice more with PBS; at this point certain cell pellets were visibly stained blue with Cy5 in a NahK- and GlcNAz-dependent manner. Cells were resuspended to a volume equal to the initial culture volume in PBS and the fluorescence of Cy5 was measured by plate reader (λ_ex_ = 610 nm, 30 nm slit width; λ_em_ = 675 nm, 50 nm slit width).

#### Nucleotide sugar metabolite extracts

LB cultures of *E. coli* BW25113 pBAD(*nahK*) (10 mL) containing antibiotics, L-arabinose (50 mg/L), and GlcNAz (1 mM) were grown at 37 °C with shaking (180 rpm) until an OD_600_ ≈ 1.0 (2-3 hours), then harvested by centrifugation (3.3 rcf; 30 minutes). Pellets were washed once in volatile ammonium bicarbonate buffer (50 mM, pH 8) to remove spent media components and resuspended in buffer (2 mL) for lysis via sonication. Lysate was cleared by centrifugation (21 k rcf; 10 minutes) and the supernatant was applied to a 3 kDa MWCO spin filter (Amicon Ultra-0.5 UFC5003) to remove larger particulates. The initial filtrate and several successive washes of the retentate with water were collected, pooled, and lyophilized to remove volatile buffer components. The lyophilized material was then reconstituted in water (200 µL) and stored at −80 °C until needed.

For those samples treated with acid to hydrolyze the carbon-phosphate glycosidic bond and release the free sugar, reconstituted samples were incubated with HCl (50 mM) and boiled for 2 minutes, then neutralized with 50 mM NaOH before fluorophore labelling. To generate the fluorophore-conjugated azide species, samples were incubated overnight with DBCO-Cy3 or DBCO-TAMRA dye (10 μM; 37 °C) before mixing 1:1 with 2× loading buffer (5 µL samples are sufficient for FACE analysis).

#### Peptidoglycan isolation and analysis

Peptidoglycan sacculi were isolated from *E. coli* BW25113 pBAD(*nahK*) cell cultures following an established protocol^3^. LB cultures (1 L) with added ampicillin (100 mg/L), arabinose (20 mg/L), and GlcNAz (0, 0.15, or 1 mM) were grown at 37 °C with shaking (180 rpm) to late log phase (OD_600_ ≈ 1.5, ∼6 hours). Cells were harvested by centrifugation (4.4 k rcf; 30 minutes) and washed once with PBS. Cell pellets were resuspended in PGN buffer (25 mM sodium phosphate pH 6.0; 10 mL) and added dropwise to boiling PGN buffer + 8% SDS (10 mL). The mixture was boiled for an additional 30 minutes to ensure maximal recovery of peptidoglycan sacculi. After removing the mixture from heat and letting it cool to room temperature, the insoluble sacculi were pelleted by centrifugation (20 k rcf; 30 minutes). The supernatant containing soluble cellular debris was carefully decanted off, and the off-white pellet was washed by resuspending with vortexing in PGN buffer (20 mL) and pelleting (20 k rcf; 30 minutes). The washes were continued until the solution stopped foaming on agitation, indicating the SDS had been removed; typically a total of 4 washes were required. The now glassy pellet was resuspended in PGN buffer (1 mL) and α-amylase (100 µg) was added. The suspension was incubated at 37 °C with turning for 1 hour, and then proteinase K (200 µg) was added and the incubation continued overnight. To remove the added enzymes, the reaction mixture was added dropwise to boiling PGN buffer + 4% SDS (10 mL) and boiled for an additional 30 minutes. After removing the mixture from heat and letting it cool to room temperature, the insoluble sacculi were pelleted by centrifugation (20 k rcf for 30 minutes). The supernatant was decanted, and the pellet was washed in an identical manner as before, until SDS was removed; typically requiring 2 washes with PGN buffer. The now purified sacculi (yield of several dozen milligrams of wet material per litre of culture) were resuspended in PGN buffer (∼4 mg/mL) and stored at 20 °C.

Digestion of the whole sacculi was done by overnight incubation with lysozyme (40 mg/mL; 37 °C) with turning. To exclude larger insoluble material, the reaction mixture was filtered through a 3 kDa MWCO spin filter (Amicon Ultra-0.5 UFC5003). The muropeptide composition of the filtrate was confirmed by ESI-MS. For FACE analysis, both whole and digested sacculi were incubated with DBCO-Cy5 dye (10 μM; 37 °C; 1 hour) before mixing 1:1 with 2× loading buffer.

#### Lipopolysaccharide isolation and analysis

Rough-type lipopolysaccharides were isolated from *E. coli* cell cultures following an established protocol^4^. Prewarmed LB cultures (50 mL) with ampicillin (100 mg/L), arabinose (20 mg/L), and GlcNAz (0-1 mM) were inoculated with starter cultures of *E. coli* containing the pBAD(*nahK*) plasmid: the wild-type BW25113 or the ECA-deficient strain AB113 *rfe*::Tn10. Both strains used are K-12 derived strains of *E. coli*, which is known to produce rough-type lipopolysaccharide. Cultures were grown overnight (180 rpm; 37 °C) and harvested by centrifugation (4.4 k rcf; 30 minutes). Cell pellets were washed with water, then ethanol, then twice with diethyl ether. After allowing ether to evaporate in an open-capped tube (rt; 60 minutes), cells were further dried by overnight lyophilization. The chalky cell pellet (25-50 mg) was resuspended in 2:5:8 aqueous phenol/chloroform/pentanes (v/v; 1 mL per 10 mg dried cells). All subsequent resuspensions were aided by water bath sonication (5 minutes) and breaking up larger clumps with a glass stir rod. The resulting suspension was left to incubate (60 °C, 15 minutes) before centrifugation (4.4 k rcf; 30 minutes). The supernatant was collected, and the above extraction was repeated twice more on the insoluble pellet. All three supernatants were pooled and filtered through filter paper, then volatile solvents were evaporated under a stream of air (2 hours). Acetone (2 volumes) was added and the solution was chilled overnight (−20 °C) to precipitate LPS. The pellet was rinsed with ice-cold acetone and collected by centrifugation (4.4 k rcf; 30 minutes), then dried under a stream of air. To remove associated proteins, the pellet was reconstituted in a minimal amount of water and treated with Proteinase K (0.5 mg/mL; 60°C; 90 minutes). The liberated lipopolysaccharides were sedimented by ultracentrifugation (100 k rcf; 2 hours) and lyophilized overnight. Samples were taken up in water and stored at 20 °C until used.

Densitometric analysis of silver-stained gels was used to normalize sample concentrations for gel analysis. Samples were incubated with DBCO-Cy5 (250 μM; 37 °C; 3 hours) before mixing 1:1 with 2× SDS sample loading buffer (120 mM Tris pH 6.8, 4% SDS (w/v), 5% (v/v) β-mercaptoethanol, 20% (v/v) glycerol) and loaded onto a 15% SDS-PAGE gel (10-30 μL). Fluorescence images of gels were acquired before removal of the gels from between their glass plates, using a G:BOX Chemi XT4 (Syngene) imager. Gels were then silver-stained using a ProteoSilver™ Silver Stain Kit (Sigma) and imaged.

#### RP-LC-MS analysis of ECA intermediates

Chemically competent ECA deletion mutants (Δ*wecA* and Δ*wecG* in MG1655 background) were transformed with *nahK* wildtype (*nahK*_WT_) and mutant (*nahK*_mut_) plasmids. A single colony of ECA mutant strains (Δ*wecA* and Δ*wecG*) carrying pBAD(*nahK*) plasmids were grown in LB media (5 mL) supplemented with arabinose (50 mg/L) and GlcNAz (150 µM). Control cells were treated identically without GlcNAz in the media. Cells were cultured with shaking at 37 °C overnight.

Cells grown overnight were pelleted (5 k rcf; 5 min) and washed with 0.9 % sodium chloride and suspended in autoclaved distilled water (700 µL) then transferred to a glass tube with 2:1 (v/v) methanol/chloroform (3 mL). These suspended cells in glass tubes were incubated at room temperature (20 min) with intermittent vortexing. Samples were transferred to a CentriVap without vacuum (20 min). Soluble supernatant was transferred to new glass tubes and were placed at −80 °C until a slurry formed and tubes were moved back to CentriVap to dry under vacuum. The vacuum dried crude cell lysate was resuspended in 1:3 n-propanol/1 % ammonium hydroxide solution (200 µL) and resuspended cells could be stored at −20 °C for one week.

Cells resuspended in 1:3 *n*-propanol/1 % ammonium hydroxide solution (v/v) were sonicated in a water bath (3 min) and spun at 14.1 k rcf (5 min). Clarified lysate (20 μL) was injected and analyzed on an Agilent 1260 Infinity II system equipped with a single quadrupole electrospray ionization MS detector using a gradient method where the gradient was 25-75 % n-propanol over 12 min with 0.1 % ammonium hydroxide as the aqueous component. Analysis was performed on a Waters BEH C18 column. SIM ions detected were 845.7 for BP, 1128.7 for BPP-GlcNAc, and 1156.8 for BPP-GlcNAz.

#### PNAG dot blots

Dot blot samples were routinely diluted 5× in water before blotting. For enzymatic treatment of samples, aliquots were incubated with DspB (5 μM) or Proteinase K (1 mg/mL) for 4-16 hours, and boiled for 2 minutes to stop enzyme activity.

Dot blots were conducted as previously described.^2^ Samples for dot blot analysis (5 μL) were blotted onto nitrocellulose membranes (BioTrace™ NT, Pall). The sample spots were left to dry completely (rt; 1 hour) before the blot was blocked with 5% BSA in PBS-T (0.1% Tween 20; 15 mL; rt; 1 hour), and subsequently incubated with HRP-WGA (0.5 mg/L; rt; 1 hour) by adding the lectin directly into the blocking buffer. The blot was washed three times with PBS-T (15 mL; rt; 10 minutes) and visualized using SuperSignal™ West Pico PLUS Chemiluminescent Substrate (Thermo Scientific). All steps with the blot immersed in solution were done with gentle shaking of the blot on a gel shaker.

#### Cellular PNAG extraction and gel analysis

Large scale cultures were required to collect a reasonable quantity of PNAG for analysis. Prewarmed LB cultures (500 mL) with kanamycin (0-50 mg/L), ampicillin (100 mg/L), arabinose (50 mg/L), and GlcNAz (0-1000 μM) were inoculated with starter cultures of *E. coli* containing the pBAD(*nahK*) plasmid: the wild-type BW25113 or the PNAG-overproducing strain MG1655 *csrA*::*kanB*. Cultures were grown aerobically and statically (rt; 2 days), and harvested by centrifugation (4.4 k rcf; 30 minutes). The cell pellets were washed in water (10 mL) and resuspended in buffer (1 M ammonium bicarbonate; 10 mL). The suspensions were left to sit at rt for 5 minutes, and centrifuged (4.4 k rcf; 30 minutes) to collect a supernatant containing extracted exopolysaccharides. Aliquots were taken for dot blot analysis and the remainder of the samples were dried down by lyophilization. To ensure removal of volatile buffer components, lyophilization was performed once more after reconstitution in water. The white fluffy lyophilized material was reconstituted in water and stored at −20 °C until needed.

To prepare samples for gel electrophoresis, samples were incubated with DBCO-Cy5 (10 μM; rt; 1 hour) and mixed 1:1 with 2× SDS sample loading buffer (120 mM Tris pH 6.8, 4% SDS (w/v), 5% (v/v) β-mercaptoethanol, 20% (v/v) glycerol). Gel samples (10 μL) were run on a 10% SDS-PAGE gel. For enzymatically treated samples, extracts were incubated with DspB (5 μM) or Proteinase K (1 mg/mL) for 4-16 hours, and boiled for 2 minutes to stop enzyme activity before mixing with loading buffer.

To reduce oligosaccharides for tandem mass spectrometry analysis, DspB-digested samples were treated with sodium borohydride (1 mg/mL) in ammonium hydroxide solution (1 M) overnight (rt), and neutralized with glacial acetic acid (1/30^th^ reaction volume). Reactions were cleaned up via graphitized carbon chromatography (see *In vitro* PgaCD reactions).

#### Cell-associated PNAG binding assay

PNAG binding assays were conducted by modification of an established protocol.^5^ Prewarmed LB media with ampicillin (100 mg/L), kanamycin (50 mg/L), arabinose (50 mg/L), and GlcNAz (0-1 mM) was inoculated with starter cultures of *E. coli* MG1655 *csrA*::*kanB* pBAD(h). Cultures were raised to mid-log phase (OD_600_ = 0.5-1.0; 37 °C; 180 rpm; 3-4 hours), harvested (6.5 k rcf, 30 minutes) and washed once with PBS. Cells were resuspended in PBS to an OD_600_ = 5.0 and incubated with azide-reactive DBCO-Cy5 (10 μM; rt; 1 hour). Cells were then washed twice with PBS and transferred to assay buffer (50 mM sodium phosphate pH 5.8, 0.5% (w/v) BSA, made up in PBS pH 5.8), and cell suspensions (0.5 mL) were applied to 0.45 μm spin filter microcentrifuge tubes (CLS8163, Corning Costar). All centrifugation steps with filter tubes were performed at 4 °C and 4 k rcf. Cell pellets were spun down in filter tubes, resuspended in buffer containing GFP-DspB^E184Q^ (0.1 μM; 150 μL), and incubated to label PNAG (rt; 10 min). Samples were filtered and the filtrate collected, and cells were resuspended in buffer without additives (150 μL) and incubated to wash the sample (rt; 10 min). In the final step, samples were were filtered and the filtrate collected, cells were resuspended in assay buffer containing active DspB enzyme (50 μM; 150 μL) and incubated to hydrolyze PNAG (rt; 60 min). After a final filtration with filtrates collected, GFP fluorescence (λ_ex_ = 470 nm, 15 nm slit width; λ_em_ = 515 nm, 20 nm slit width) and Cy5 fluorescence (λ_ex_ = 610 nm, 30 nm slit width; λ_em_ = 675 nm, 50 nm slit width) of the collected filtrates were measured by plate reader.

To account for background autofluorescence of cells, values were corrected using a sample with *E. coli* MG1655 *csrA*::*kanB* pBAD(*nahK*) cells treated by a modified protocol without any added DBCO-Cy5 or GFP-DspB^E184Q^. All fluorescence values were reported as relative values: relative to the initial labelling concentration of GFP-DspB^E184Q^ (0.1 μM), or the fold increase in Cy5 signal relative to the blank sample for GFP and Cy5 fluorescence respectively.

#### Crystal violet biofilm assay

Starter cultures of *E. coli* MG1655 *csrA*::*kanB* pBAD(*nahK*) were used to inoculate fresh LB media with ampicillin (100 mg/L), kanamycin (50 mg/L), arabinose (20 mg/L), and GlcNAz derivative (0-1 mM). Inoculated media was plated into the wells of a clear 96-well flat-bottomed microplate (200 μL) and incubated (rt; 24 hour) to allow biofilm growth. Spent media and planktonic cells were removed from the wells and the plate was washed once with water. Adherent biomass was stained with crystal violet stain (0.1% w/v; 200 μL; 30 minutes). The stain was removed from the wells and the plate was washed once with water. The wells were left to air dry overnight and the stain was solubilized in 30% acetic acid (200 μL; 30 min). The concentrated dye solutions were diluted into a new microplate with 30% acetic acid to lower the optical density, and the absorbance at 550 nm was measured by plate reader.

#### *In vitro* PgaCD reactions

Reactions with either purified PgaCD_m_ or empty BL21 DE3 membranes (88 mg/mL) were carried out in reaction buffer (50 mM HEPES pH 7.5, 300 mM NaCl, 5 mM MgCl_2_, 5% glycerol) using a mixture of UDP-GlcNAc and UDP-GlcNAz (25 mM total) as sugar donor substrates. Reactions were incubated 24 hours at 37 °C and aliquots were taken for dot blot analysis. PNAG polysaccharide was subsequently hydrolyzed with DspB treatment (5 μM; 100 mM sodium phosphate pH 5.8; 37 °C; 90 minutes) to afford water-soluble oligosaccharides. Reaction mixtures were boiled (2 minutes) and clarified by centrifugation (21 k rcf; 10 minutes). Clean fractions of oligosaccharide were obtained via graphitized carbon chromatography using Hypercarb™ Hypersep™ SPE cartridges (60106-301, Thermo Scientific). Samples were applied to pre-equilibrated columns and washed with 6 mL water before elution with 25% (v/v) ACN/water. Eluates were lyophilized and reconstituted in volumes of water equivalent to the initial reaction volumes, and stored at −20 °C until needed.

To label azides present in cleaned oligosaccharide, samples were incubated with DBCO-Cy5 (10 μM; rt; 1 hour). For FACE gel electrophoresis, labelled samples (20 μL) were lyophilized and reconstituted in 2× FACE loading buffer (10 μL).

**Figure S1.**
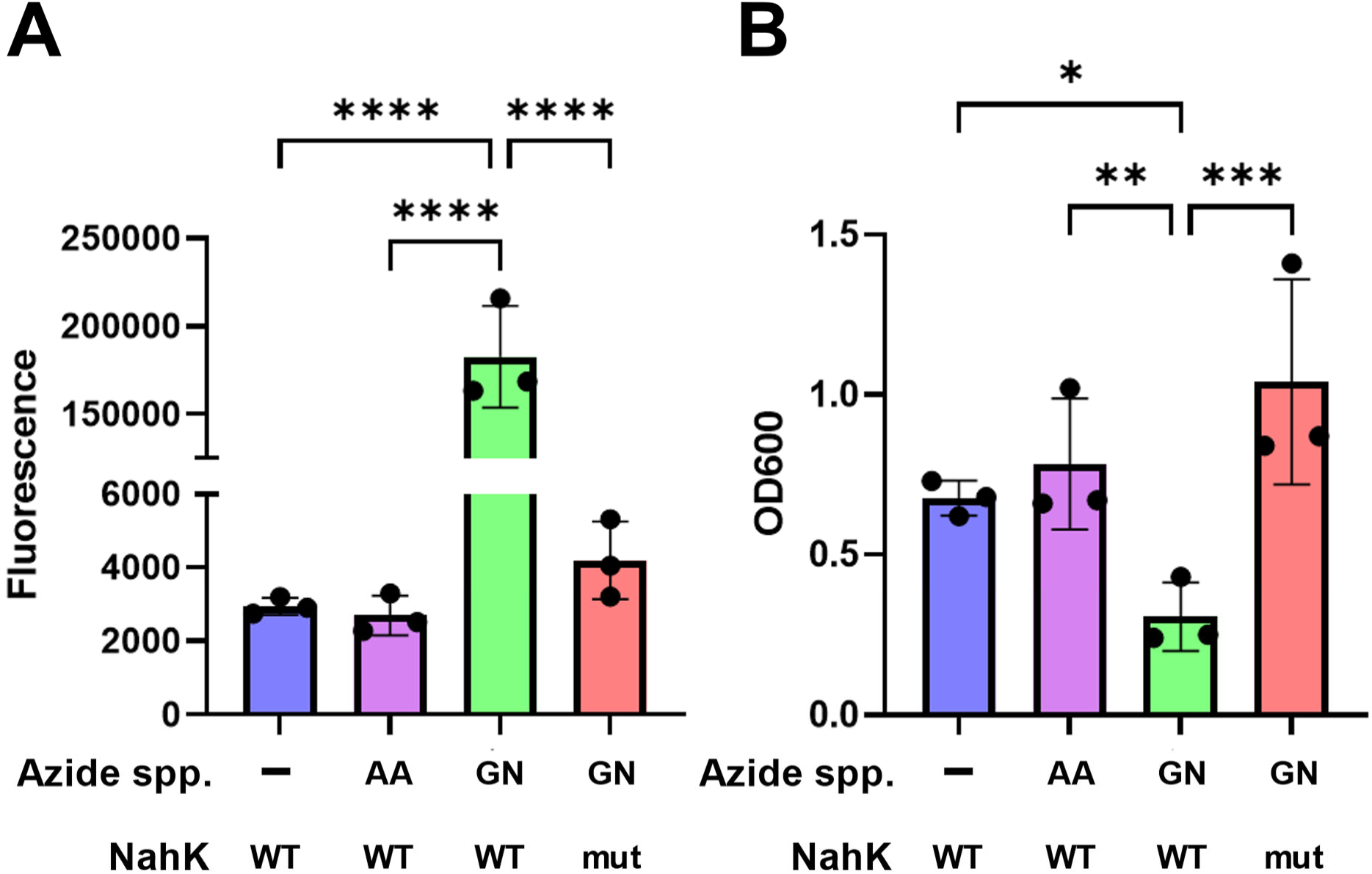
GlcNAz incorporation into whole cells. *E. coli* BW25113 pBAD(*nahK*) were grown with azide species: GlcNAz (GN; 1 mM) or AzAcOH (AA; 1 mM). Cells were incubated with DBCO-Cy5 and measured for retained Cy5 fluorescence (**A**) and cell density (**B**) after several cell washes. Error bars represent ±SD of 3 biological replicates; significance was determined using unpaired t-tests. ****P ≤ 0.0001, ***P ≤ 0.0008, **P ≤ 0.0072, *P ≤ 0.023.

**Figure S2.**
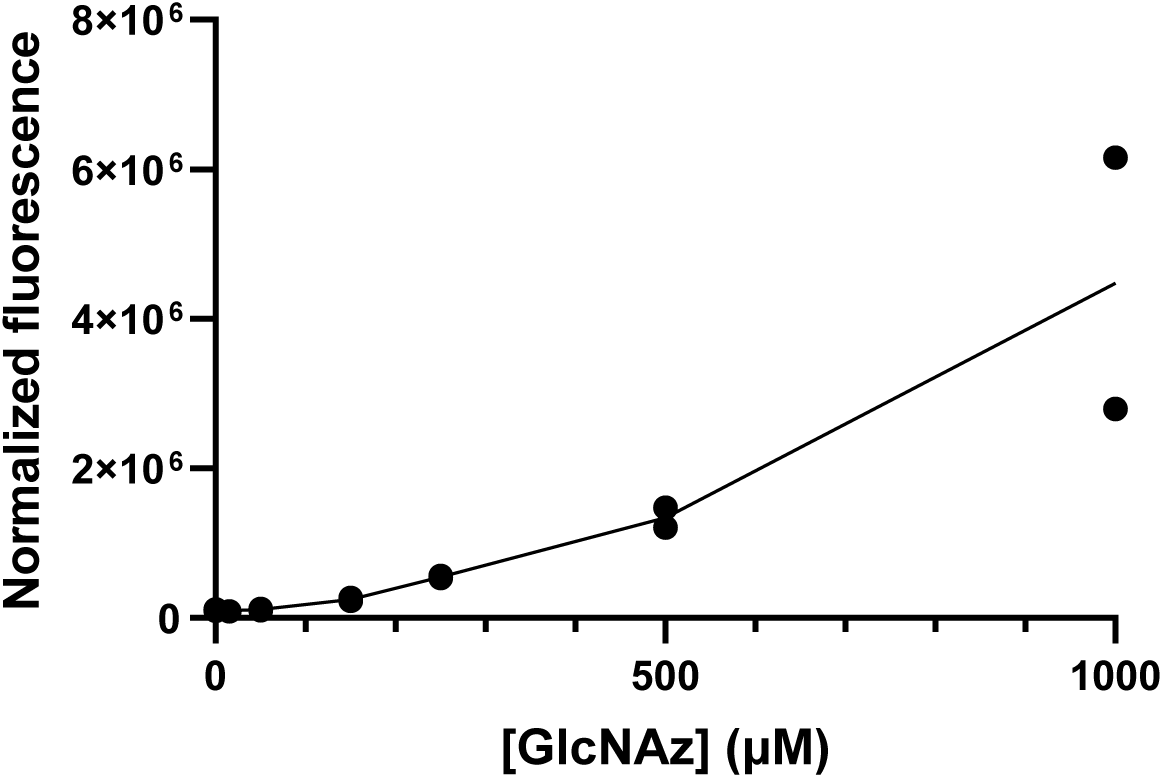
GlcNAz dose-response. *E. coli* BW25113 pBAD(nahK_WT_) cultures were grown with GlcNAz and labelled with DBCO-Cy5. Cells were washed several times to remove free dye and measured for cell-normalized Cy5 fluorescence. Data is taken from two biological replicates.

**Figure S3.**
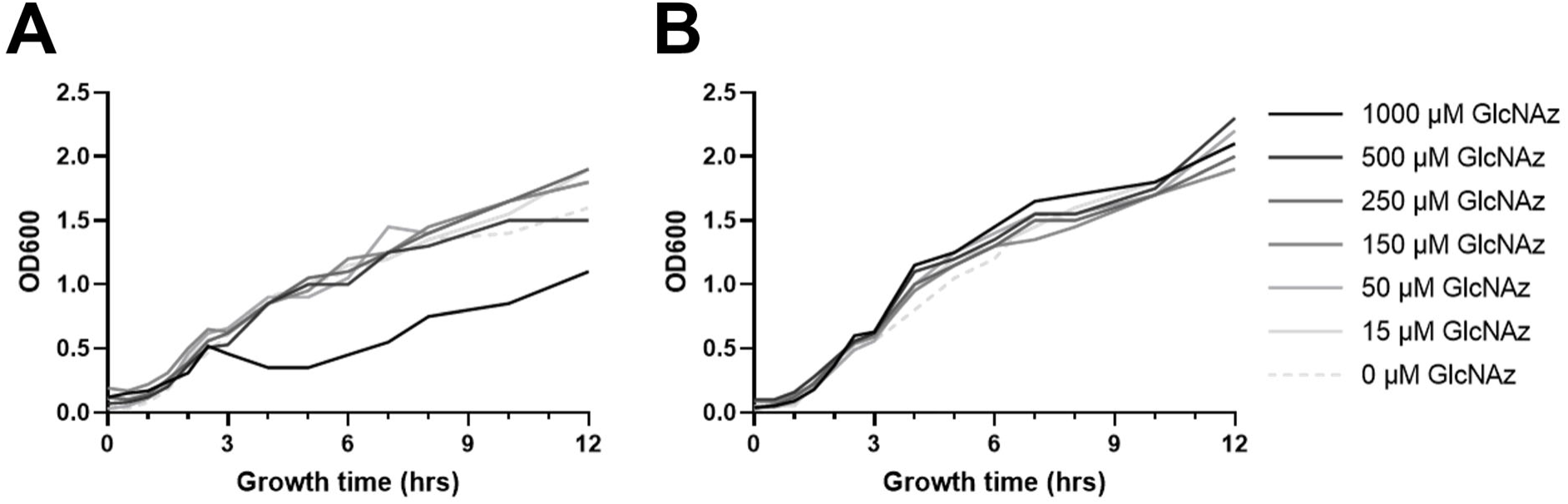
Growth curve with GlcNAz. *E. coli* BW25113 (**A**) pBAD(*nahK*_WT_) or (**B**) pBAD(*nahK*_mut_) cultures were grown with GlcNAz for several hours. Data represents the average of two biological replicates.

**Figure S4.**
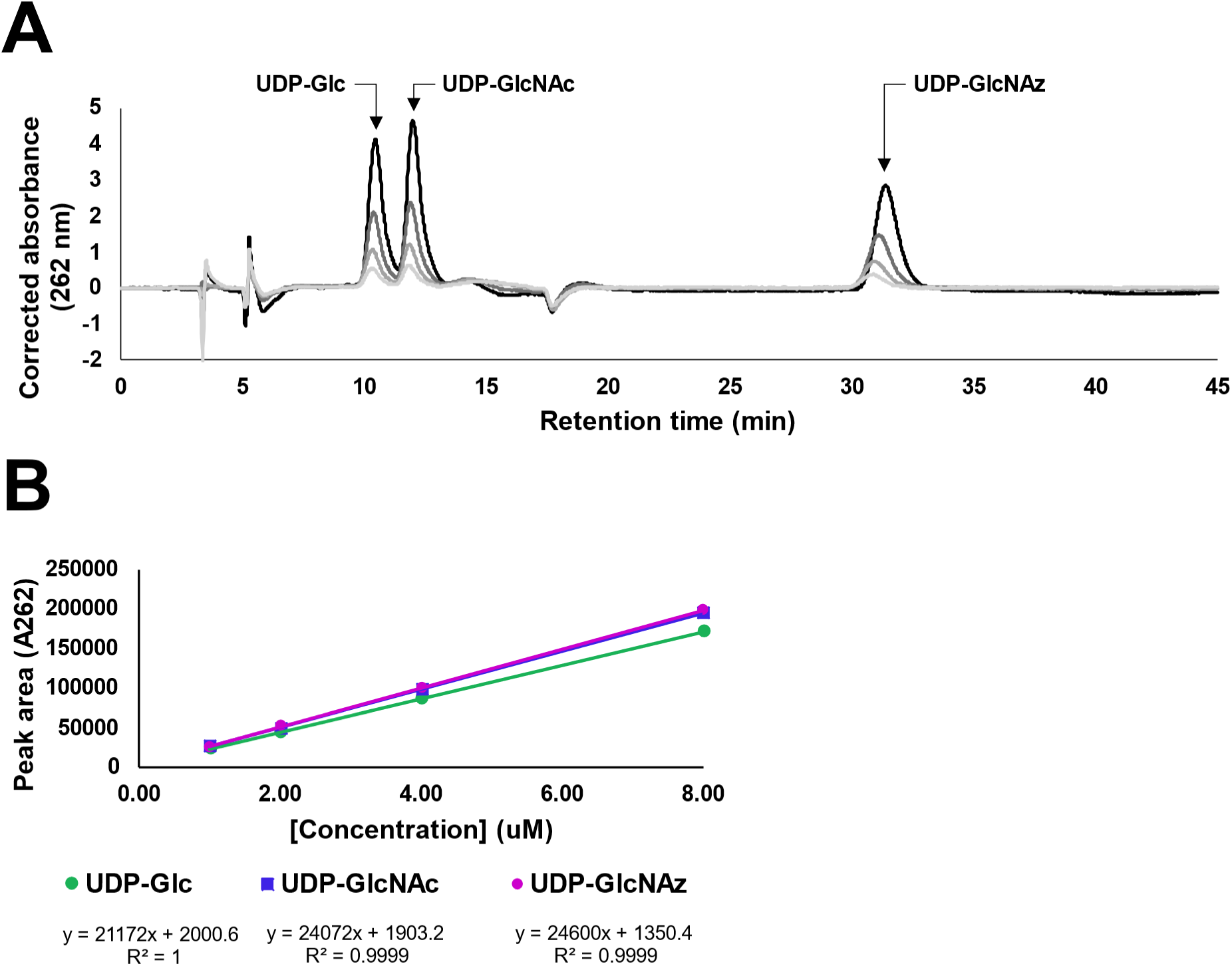
HPLC of sugar-nucleotides. (**A**) Representative ion-pairing reverse-phase HPLC trace of UDP-sugar standards at 150 mM potassium phosphate in mobile phase. (**B**) Calibration curve of sugar nucleotide concentrations derived from measuring peak areas of the preceding trace.

**Figure S5.**
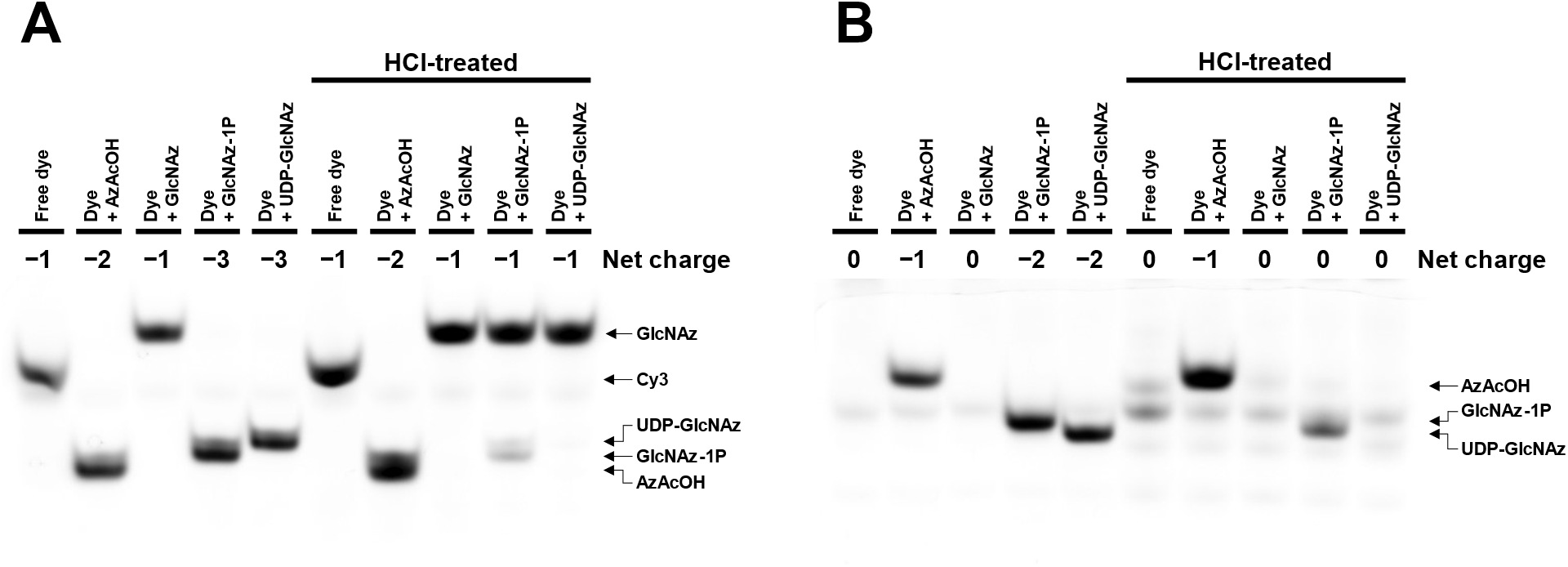
FACE gel standards. Aliquots (5 μL) of each azide-containing species (0.5 mM) were selectively treated with acid to hydrolyze sugar-phosphate bonds and incubated with 10 μM (**A**) DBCO-Cy3 or (**B**) DBCO-TAMRA. FACE gel was run and imaged for fluorescence. Arrows indicate migration distances of various tagged azide species; distances can be correlated with net charge of each species.

**Figure S6.**
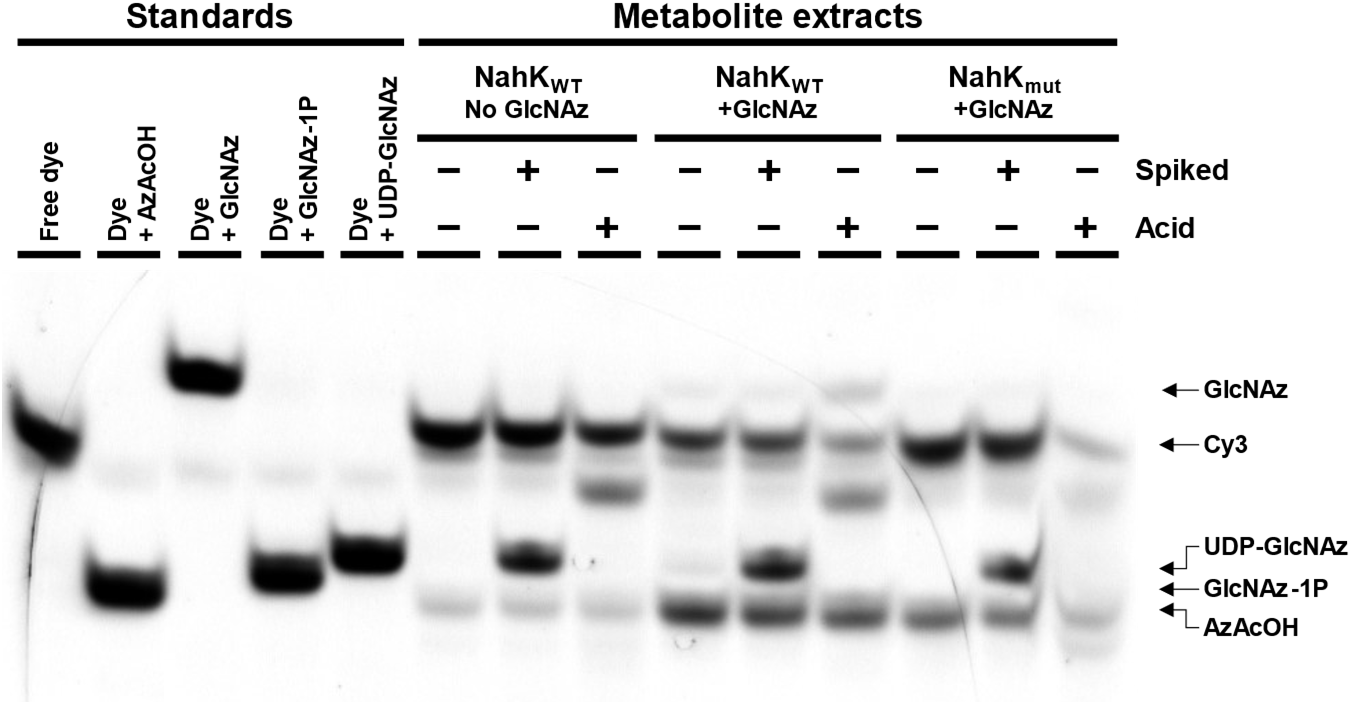
Cy3-tagged metabolite FACE gel. Nucleotide sugar extracts of GlcNAz-fed *E. coli* BW25113 pBAD(*nahK*) cells were selectively heated with acid to hydrolyze sugar phosphates (**Acid**), incubated with DBCO-Cy3 to tag azides, and spiked with UDP-GlcNAz migration standard (**Spiked**). FACE gel was run with migration standards of tagged azide species and imaged for Cy3 fluorescence. Arrows indicate migration distances of various tagged azide species.

**Figure S7.**
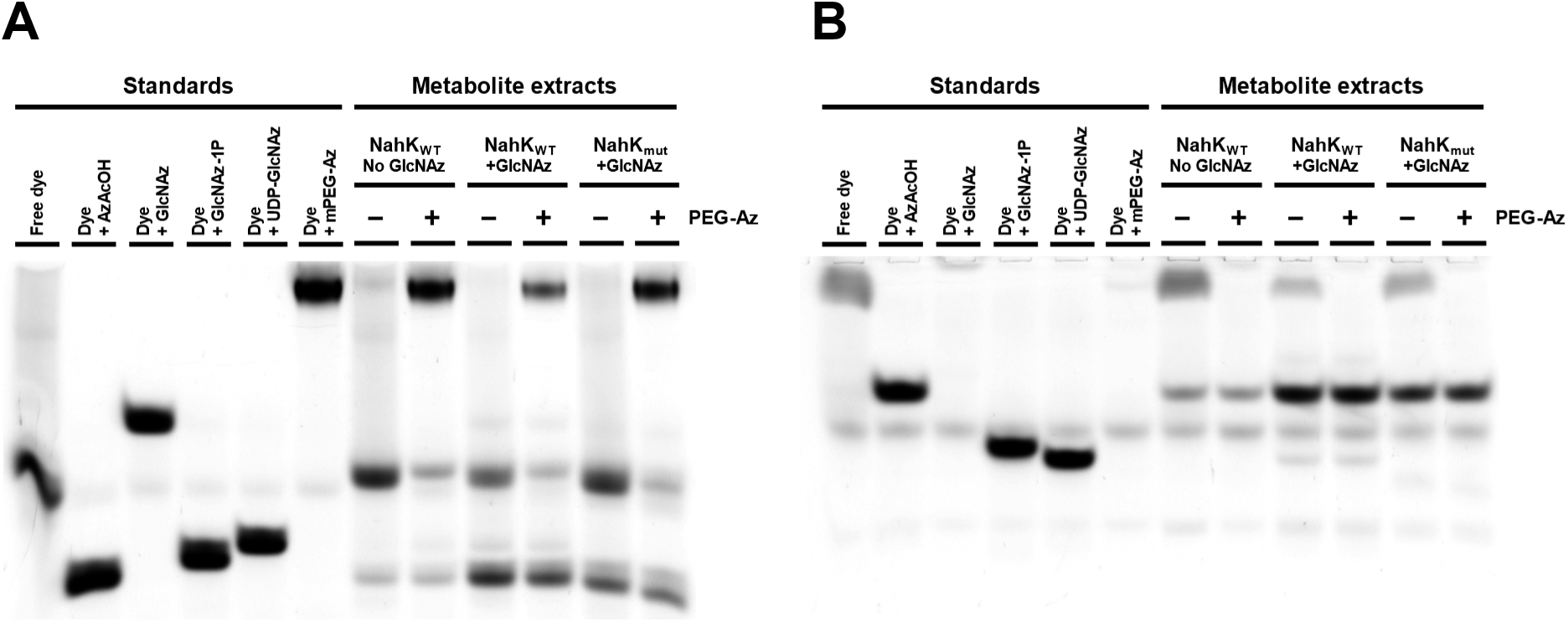
PEG-Az metabolite FACE gel. Nucleotide sugar extracts of GlcNAz-fed *E. coli* BW25113 pBAD(*nahK*) cells were incubated with (**A**) DBCO-Cy3 or (**B**) DBCO-TAMRA to tag azides, and selectively treated with PEG-Az to sequester any excess free dye. FACE gel was run with migration standards of tagged azide species and imaged for fluorescence. Arrows indicate migration distances of various tagged azide species.

**Figure S8.**
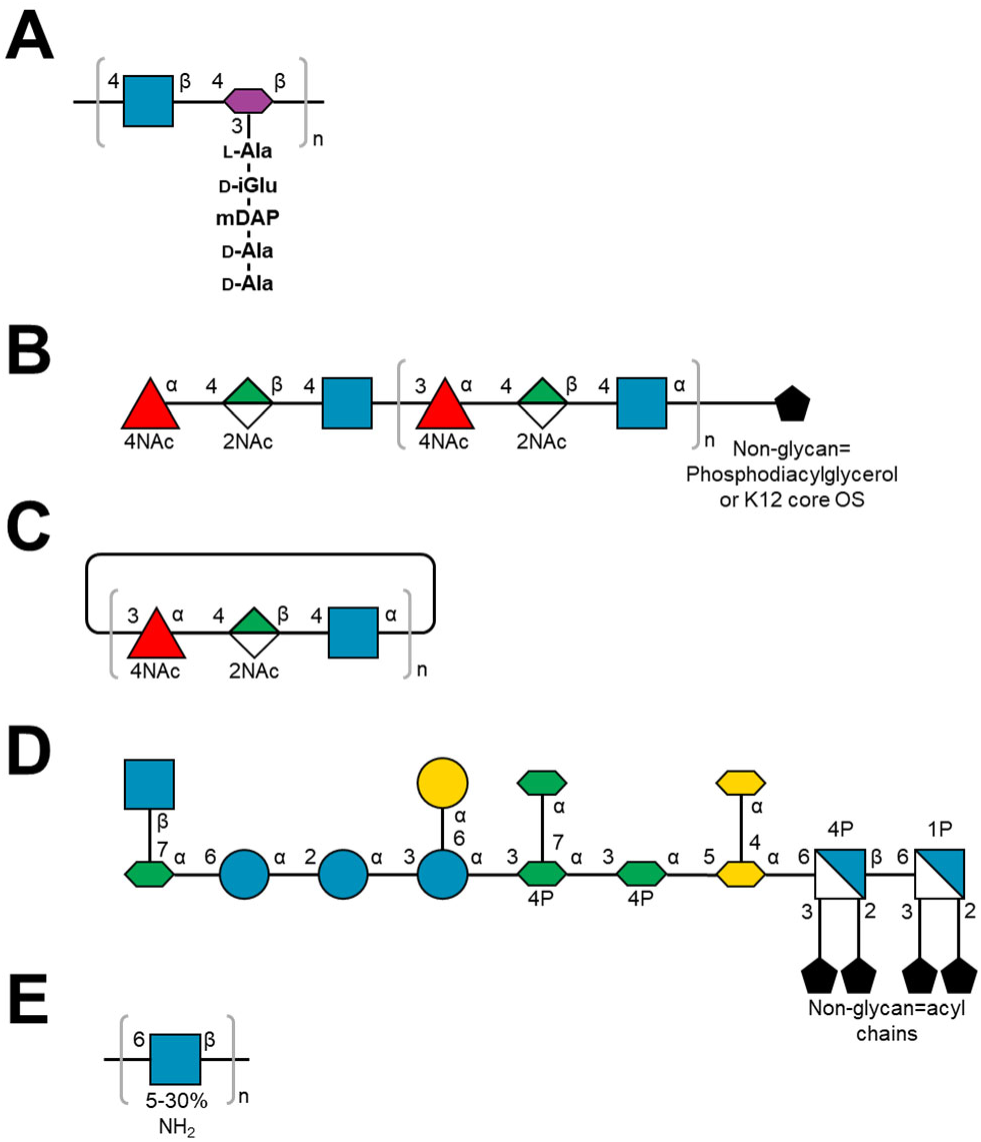
Glycan structures. (**A**) Repeating unit of *E. coli* peptidoglycan; N-terminus stem peptide is connected to MurNAc via ether linkage at lactate moiety. Peptide may lack one or both terminal D-Ala, and possess amide bond crossbridge with an adjacent peptide through mDAP or the non-terminal D-Ala. (**B**) Structure of ECA linked with diacyglycerol (ECA_PG_) or the core oligosaccharide of E. coli K12 (ECA_LPS_). (**C**) Structure of cyclic ECA; typically 4-6 repeats. (**D**) Structure of the core oligosaccharide of *E. coli* K12. (**E**) Repeating unit of PNAG; the degree of deacetylation varies depending on species and environmental conditions, and the chain length may exceed several hundred GlcNAc units. Structures are depicted using the symbol nomenclature for glycans.^62,63^

**Figure S9.**
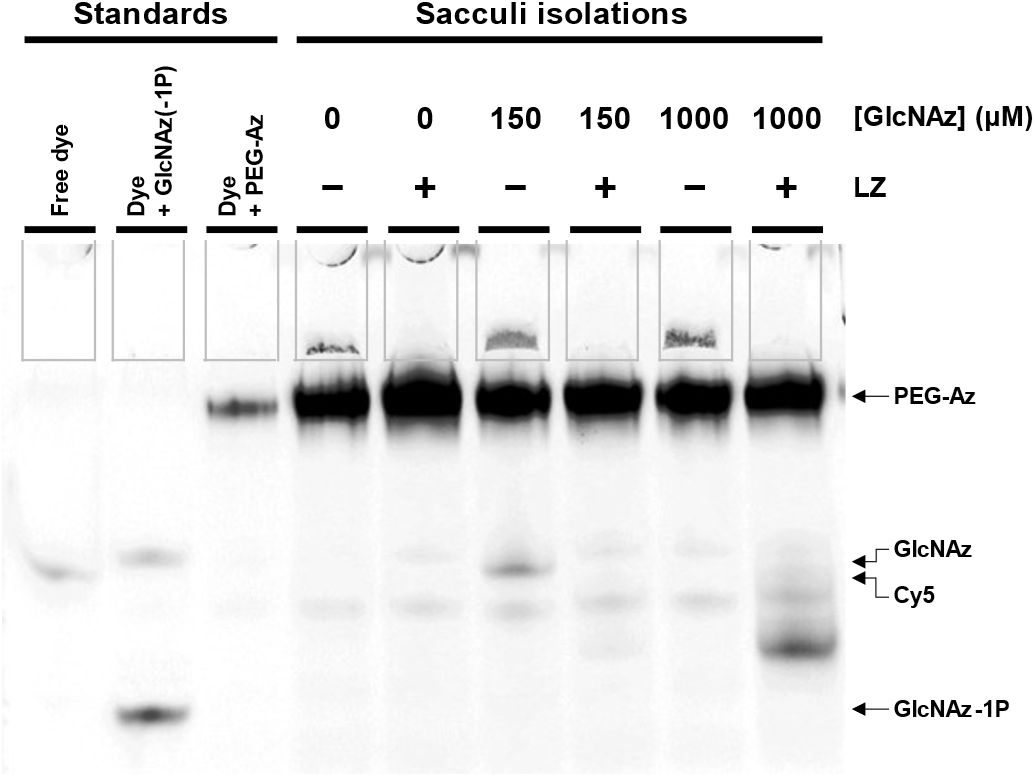
PEG-Az peptidoglycan FACE gel. Isolated sacculi from GlcNAz-fed *E. coli* BW25113 pBAD(*nahK*) cells were selectively treated with lysozyme (LZ) to generate muropeptides, incubated with DBCO-Cy5 to tag azides, and selectively treated with PEG-Az to sequester any excess free dye. FACE gel was run with migration standards of various tagged azide species (GlcNAz(−1P) standard was generated as an incomplete NahK reaction with GlcNAz) and imaged for Cy5 fluorescence. Arrows indicate migration distances of various tagged azide species. Sample wells are outlined for ease of viewing.

**Figure S10.**
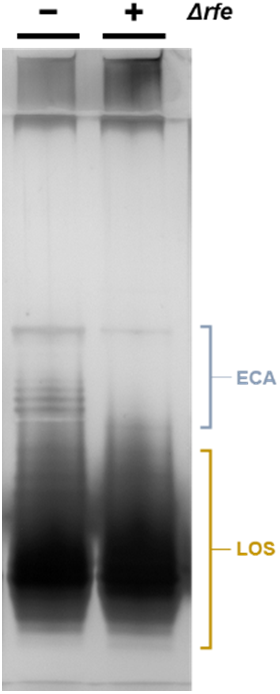
Silver-stained gel of PCP extracts. PCP extract samples from *E. coli* K12 strains analyzed by silver staining show presence of rough-type LPS (marked in brown) and lipid-linked ECA (marked in gray). Δ*rfe* strain does not display ECA bands.

**Figure S11.**
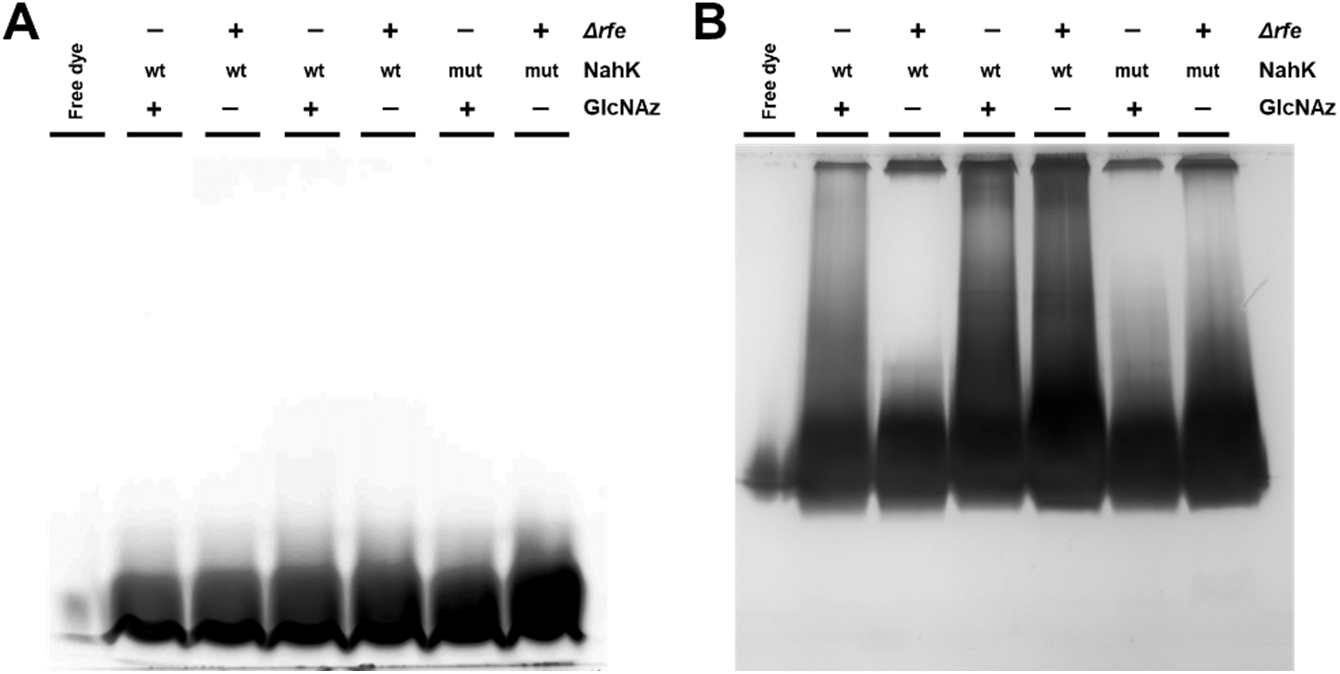
LOS/ECA SDS-PAGE gels. Isolated LOS/ECA from GlcNAz-fed *E. coli* pBAD(*nahK*) cells were incubated with DBCO-Cy5 to tag azides. PAGE gel was run with a migration standard of free dye. Efforts were made to normalize sample loading by a through a preliminary silver-stained gel. Gel was imaged for (**A**) Cy5 fluorescence and (**B**) silver-stained to detect LOS/ECA. Δ*rfe*: *rfe*::Tn10 knockout strain that is ECA^-^.

**Figure S12.**
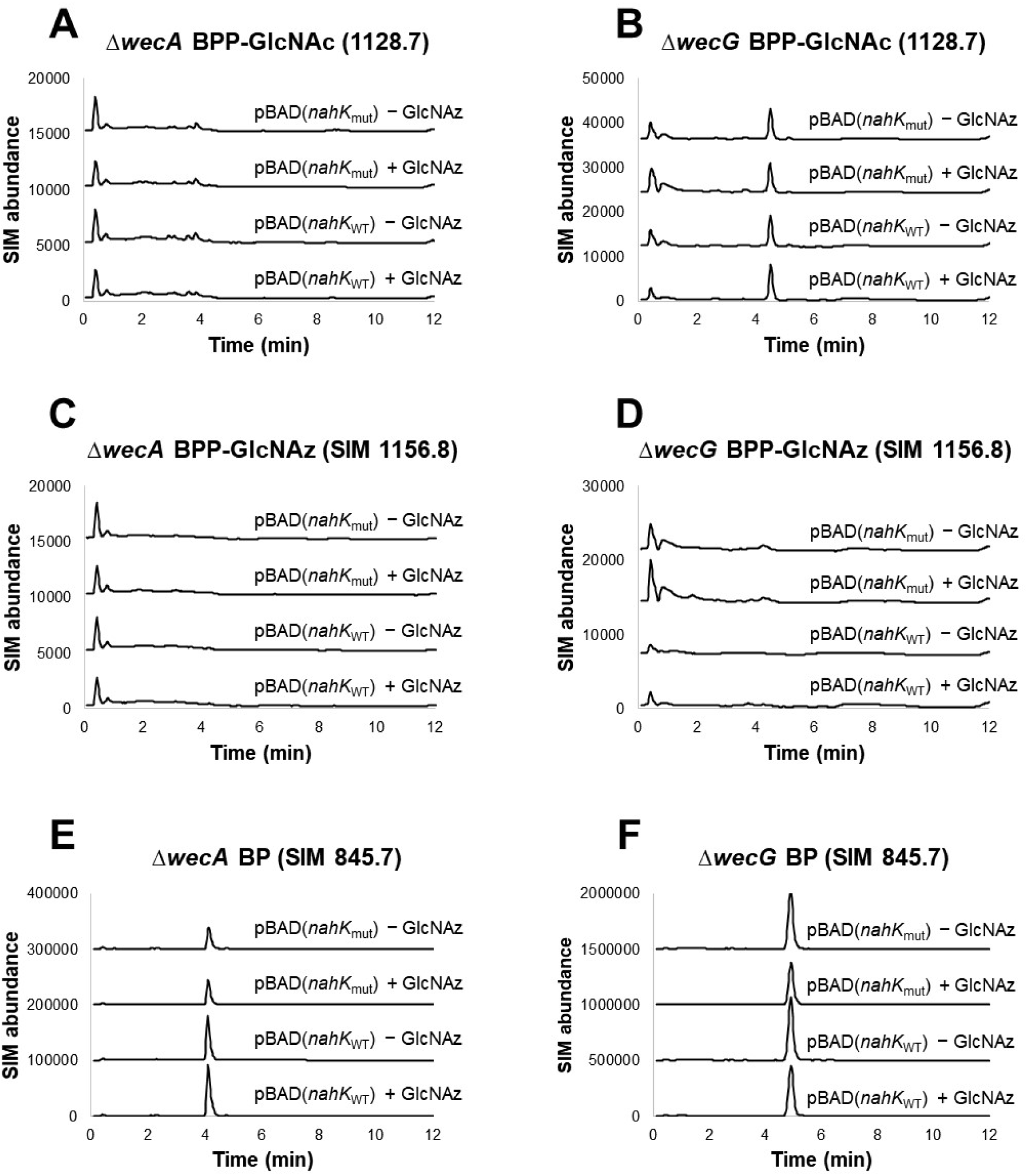
RP-LC-MS SIM of BPP-linked intermediates. Lysates from ECA knockout strains of *E. coli* MG1655 pBAD(*nahK*) grown with GlcNAz were analyzed by LC-MS, monitoring (**A**,**B**) BPP-GlcNAc, (**C**,**D**) BPP-GlcNAz, and (**E**,**F**) BP. Samples from Δ*wecA* strains (**A,C,E**) lack any ion signals consistent with any lipid-linked intermediates, and samples from Δ*wecG* strains (**B**,**D**,**F**) lack any ion signals consistent with BPP-GlcNAz.

**Figure S13.**
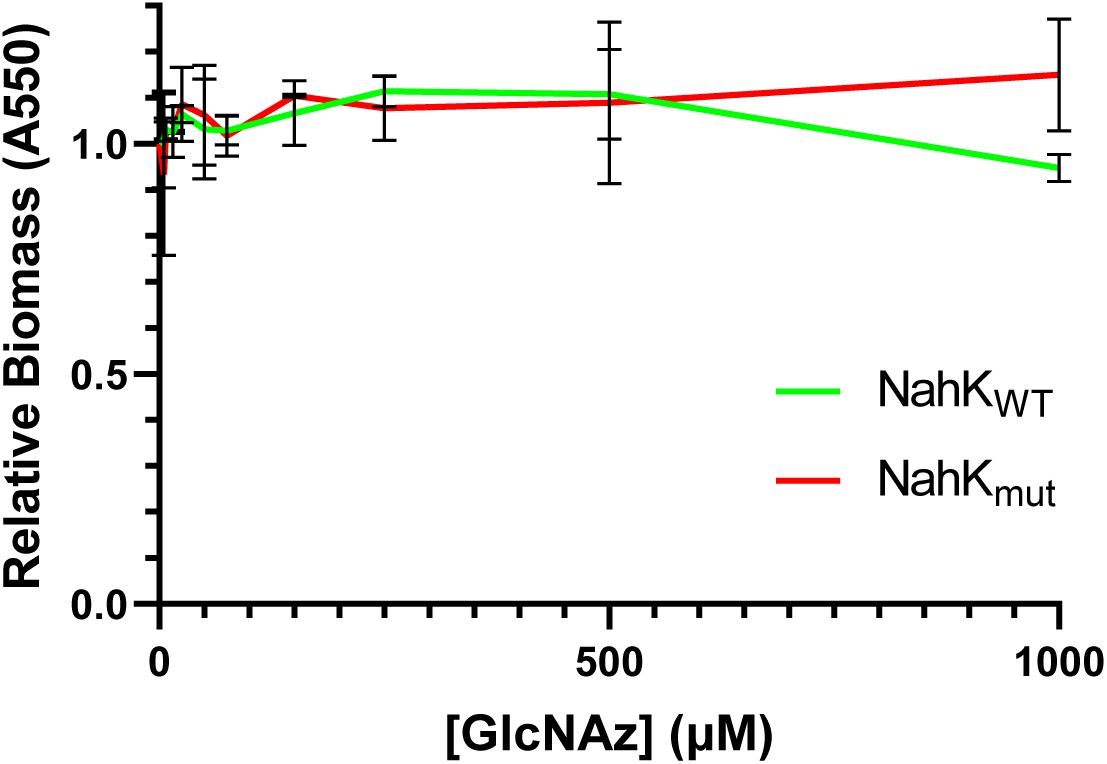
Crystal violet biofilm assay with GlcNAz. Standing cultures of *E. coli csrA*::*kanB* (**A**) pBAD(*nahK*_WT_) or (**B**) pBAD(*nahK*_mut_) cells were grown with GlcNAz to form PNAG-dependent biofilms. Adherent biomass was stained with crystal violet and quantified by absorbance at 550 nm.

**Figure S14.**
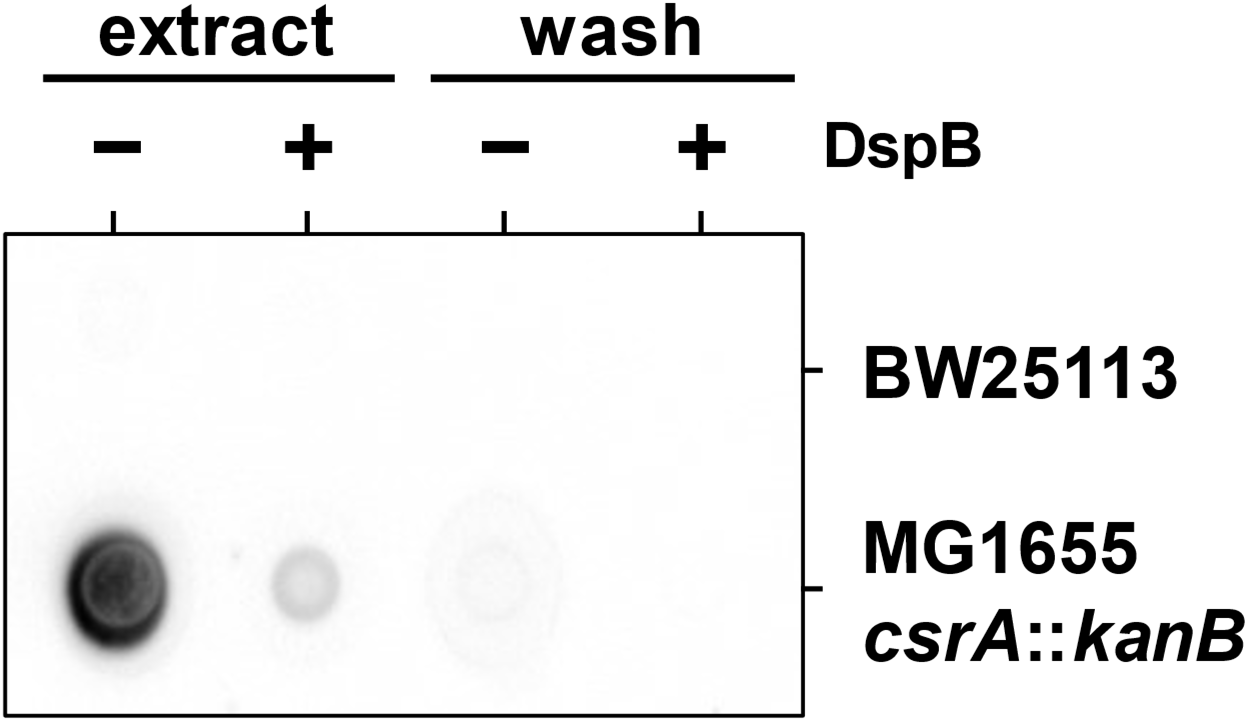
PNAG extraction dot blot. *E. coli* pBAD(*nahK*) cultures were washed with water (**wash**) and incubated in ammonium bicarbonate buffer (**extract**) to extract PNAG. Fractions were spotted on nitrocellulose membranes and probed with the GlcNAc-specific lectin fusion WGA-HRP. Select samples were pretreated with the PNAG-specific hydrolase DspB. The majority of luminescence is confined to the post-water wash extract fractions. Overproducing strains of PNAG (Δ*csrA*) generate more signal than the wildtype control strain.

**Figure S15.**
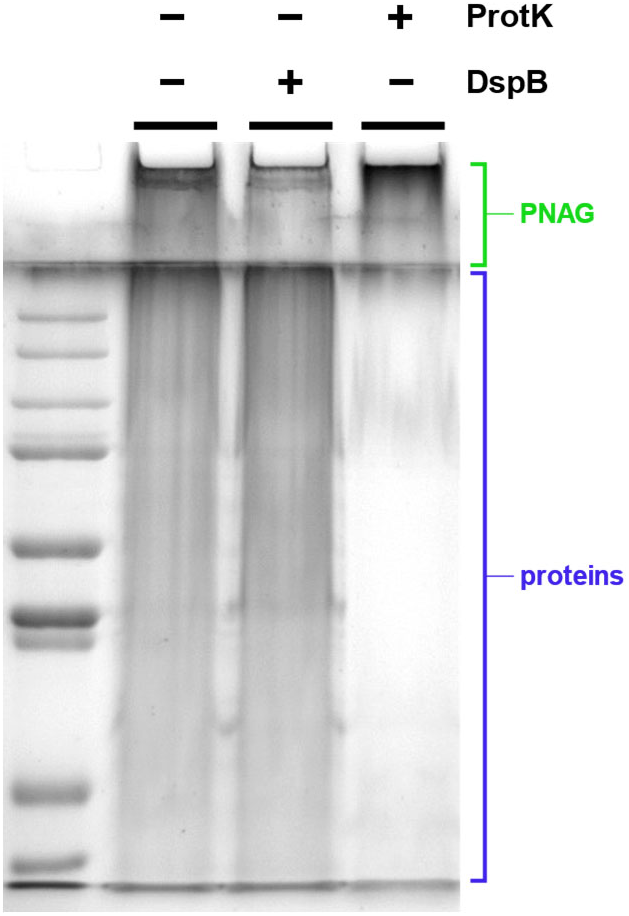
Enzymatic treatment of PNAG extracts. PNAG extracts from *E. coli* MG1655 *csrA*::*kanB* cultures were treated with DspB or ProtK and analyzed by PAGE with Coomassie Brilliant Blue staining. Compared to the control lane, the proteolyzed sample displays significantly less staining in the lower portion of the gel (proteins marked in blue) but retains some degree of staining in the upper portion, below the sample well. In contrast, the DspB-treated sample lane shows decrease in stained material in the sample well (PNAG marked in green) but retains all the protein-specific staining.

**Figure S16.**
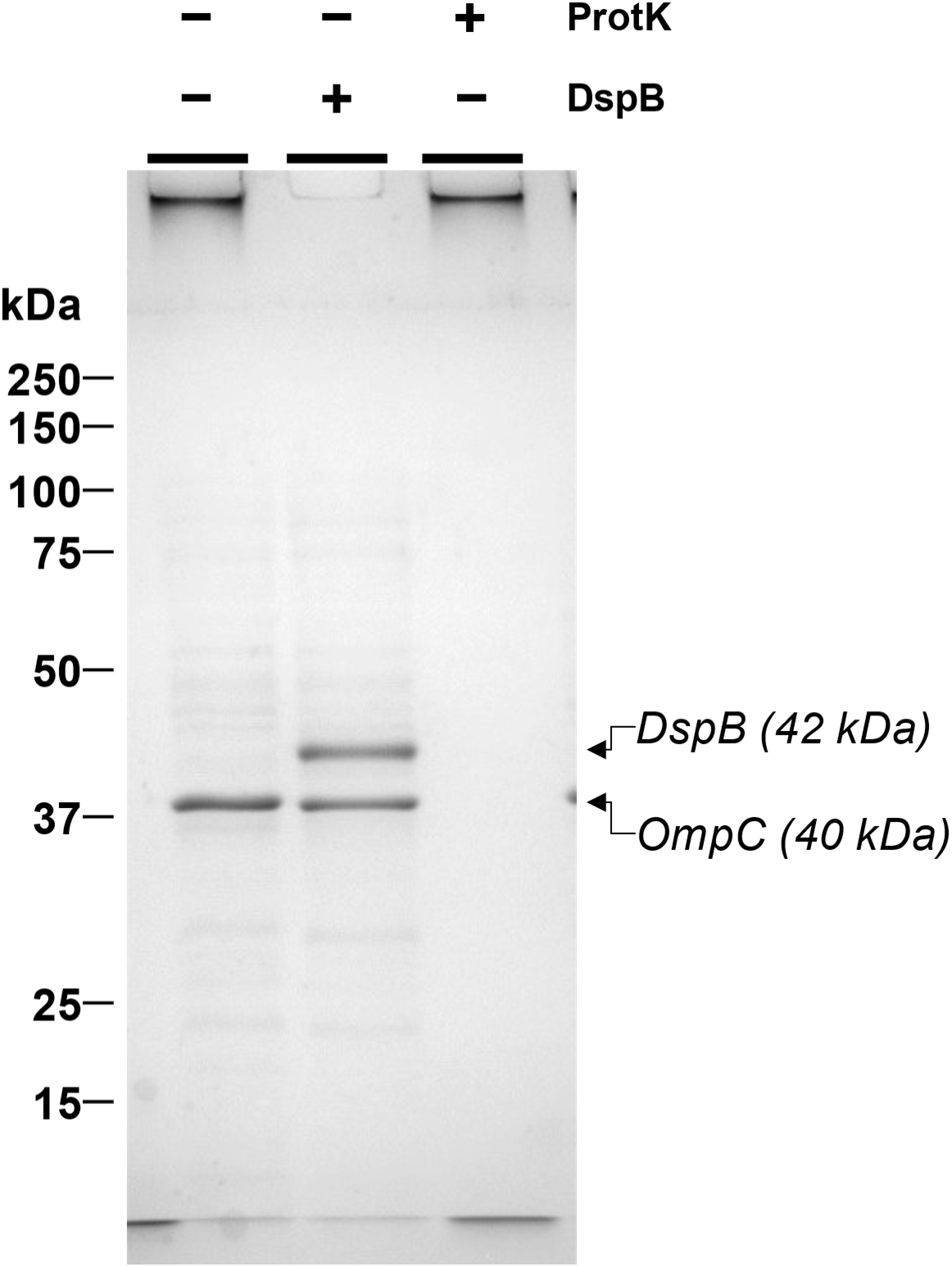
OmpC in PNAG extracts. PNAG extracts from *E. coli* MG1655 *csrA*::*kanB* cultures were analyzed by SDS-PAGE. One major protein band around 40 kDa was cut out of the gel for tryptic digest and peptide mass spectrometry fingerprinting; this protein was identified as the abundant *E. coli* outer membrane porin C (OmpC). Proteomics analysis was performed at the SPARC BioCentre at SickKids.

**Figure S17.**
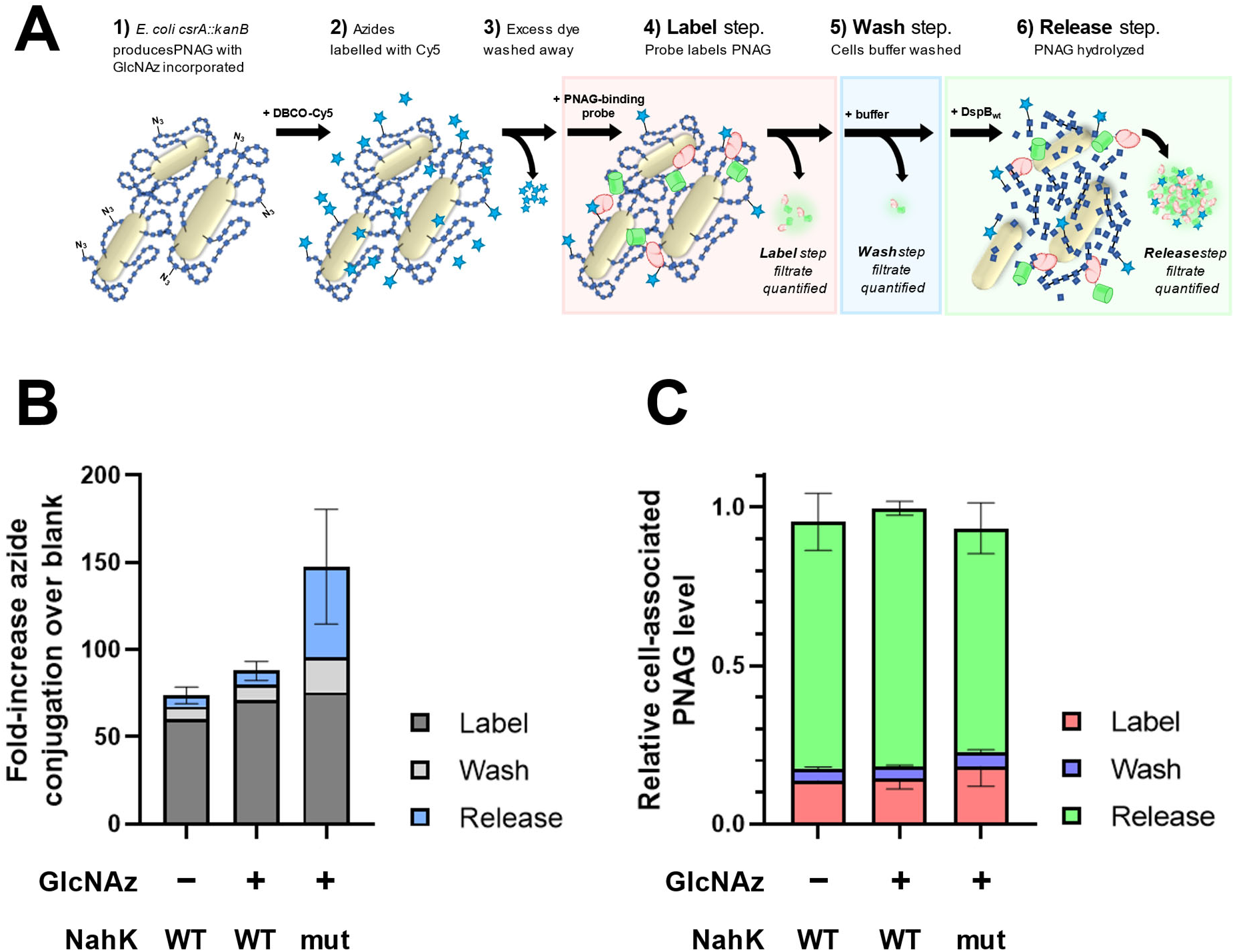
Cell-associated PNAG assay breakdown. (**A**) GlcNAz-fed *E. coli csrA*::*kanB* pBAD(*nahK*) cells were labelled with DBCO-Cy5 to tag azides, and incubated with PNAG-binding GFP-DspB^E184Q^. Cells were washed with buffer and DspB enzyme to hydrolyze PNAG. Data plotted represents (**B**) the Cy5 and (**C**) GFP signals in each collected filtrate fraction. Cy5 values are relative to a blank sample without added dye; GFP values are normalized to the fluorescence of the initial labelling solution. Error bars represent ±SD of 3 biological replicates. High Cy5 signal in **Label** fractions can be attributed to a small number of washes after incubation in the DBCO-Cy5 labelling solution; more washes decreased this signal without changing the Cy5 signal in **Release** fractions, and were therefore deemed unnecessary.

**Figure S18.**
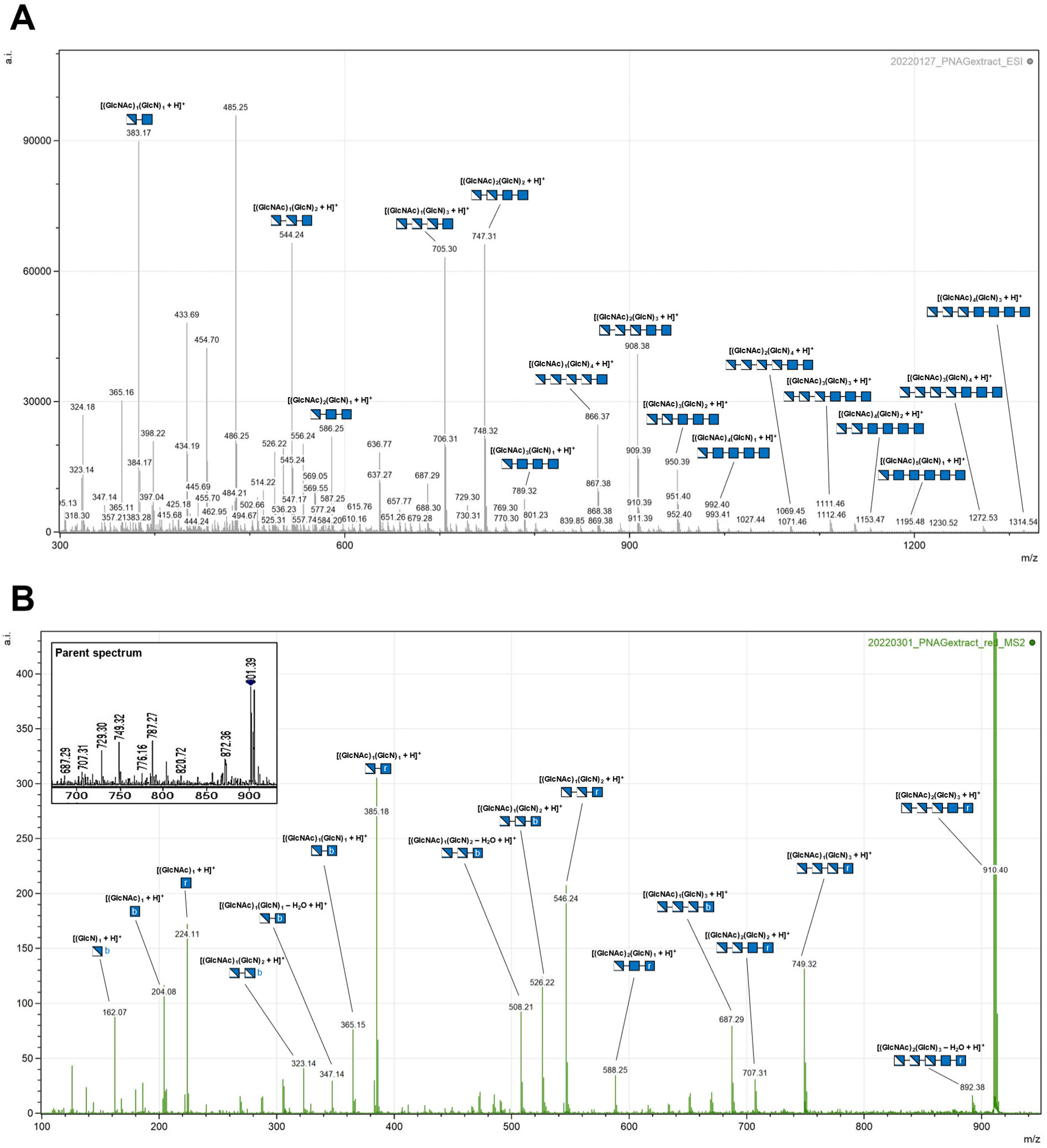
ESI-MS of PNAG extract. Deproteinized PNAG extracts from *E. coli* MG1655 *csrA*::*kanB* cultures were digested with DspB and purified via graphitized carbon chromatography. (**A**) ESI-MS analysis shows PNAG oligomers with high degrees of deacetylation. (**B**) Chemically-reduced samples were analyzed by tandem ESI-MS-MS. A pentasaccharide containing 3 positions of deacetylation was selected as the parent peak. B-ions are marked with **b**, and reducing end alditols (y-ions) are marked with **r**. The positions of deacetylation displayed in the glycan structures are for illustrative purposes only; the exact positions were not determined. It is however apparent that few intact oligomers contain GlcN at the reducing end, likely due to the known site selectivity of DspB.^64^

**Figure S19.**
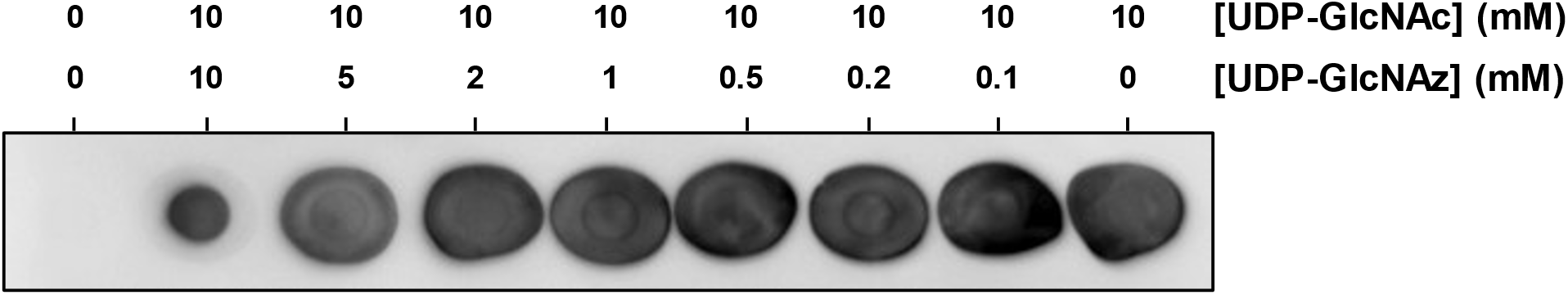
PgaCD polymerase inhibition. Dot blot assay of PgaCD reactions, holding a constant [UDP-GlcNAc] and increasing [UDP-GlcNAz], show PgaCD reaction progress decreasing as a function of UDP-GlcNAz concentration. This apparent inhibitory effect is weak, only giving significant reduction of signal when

**Figure S20.**
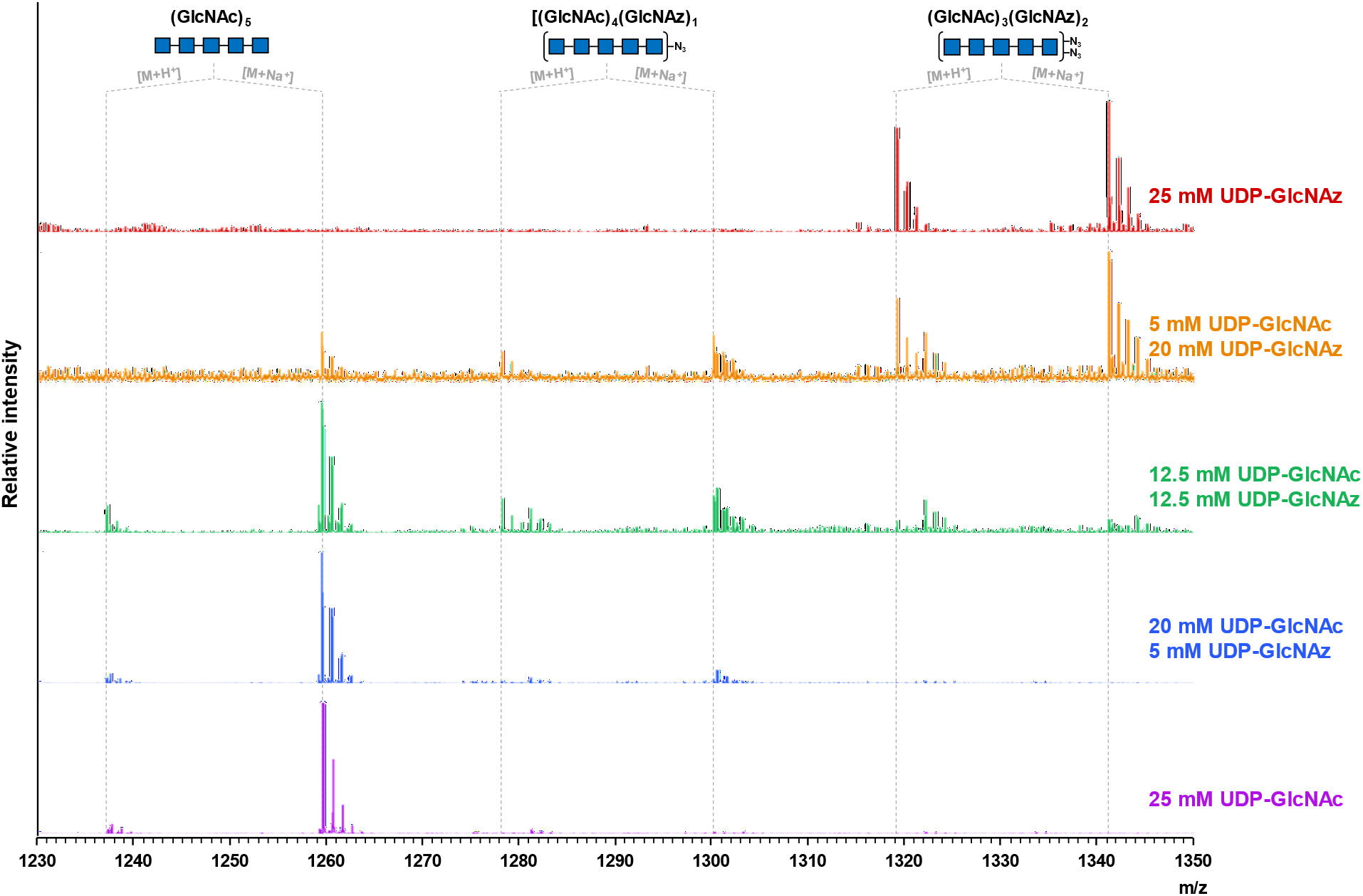
ESI-MS of PgaCD reaction. PgaCD reaction mixtures using UDP-GlcNAc and -GlcNAz as sugar donors were digested with DspB. Pentasaccharide ions containing 0, 1, or 2 GlcNAz units are displayed. Peak intensities are relative to the highest intensity signal in each respective spectrum.

**Figure S21.**
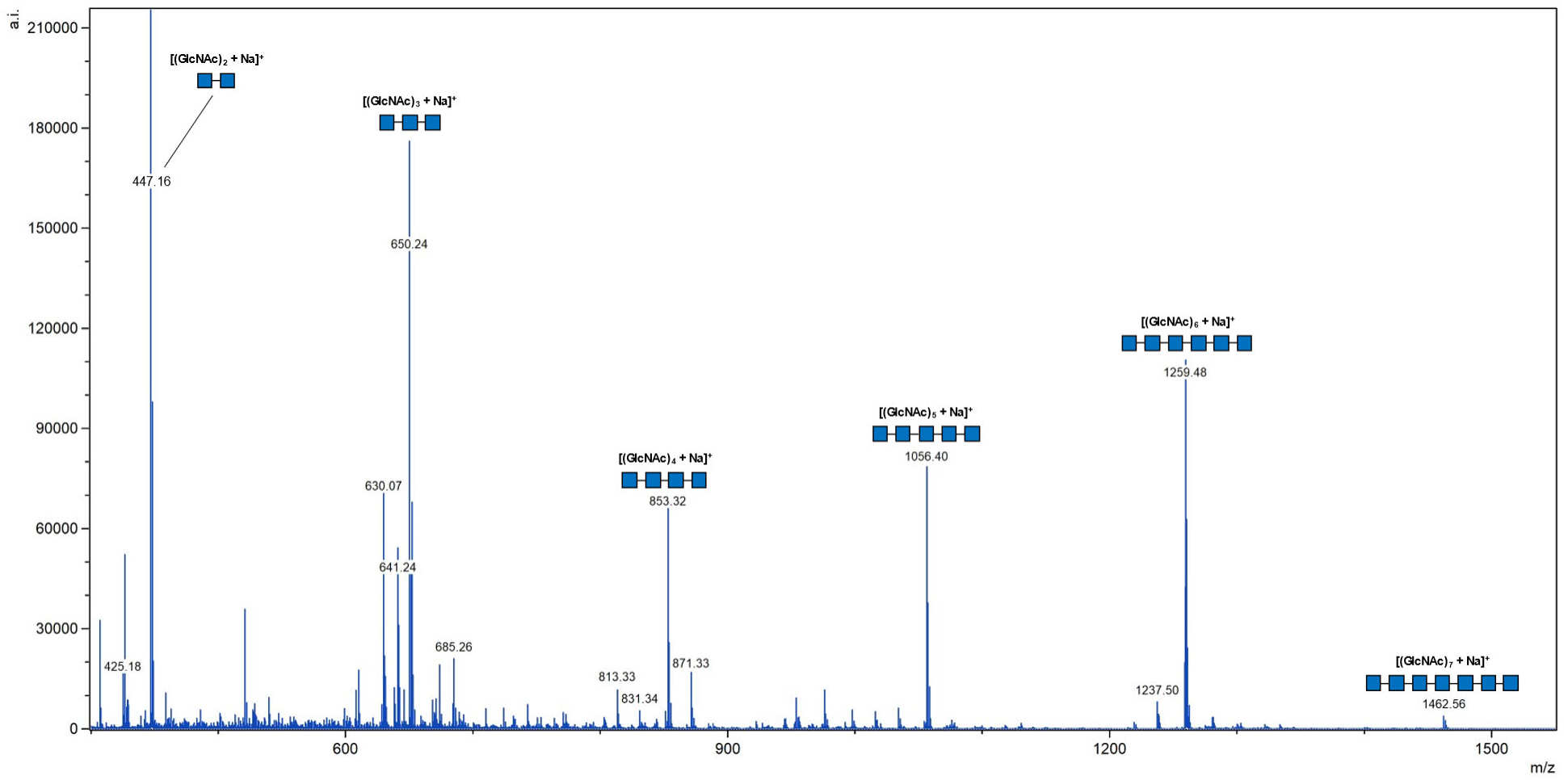
PgaCD reaction mixture using UDP-GlcNAc (25 mM) as sugar donor was digested with DspB. Dominant peak of each species is the sodium adduct [M+Na]^+^.

**Figure S22.**
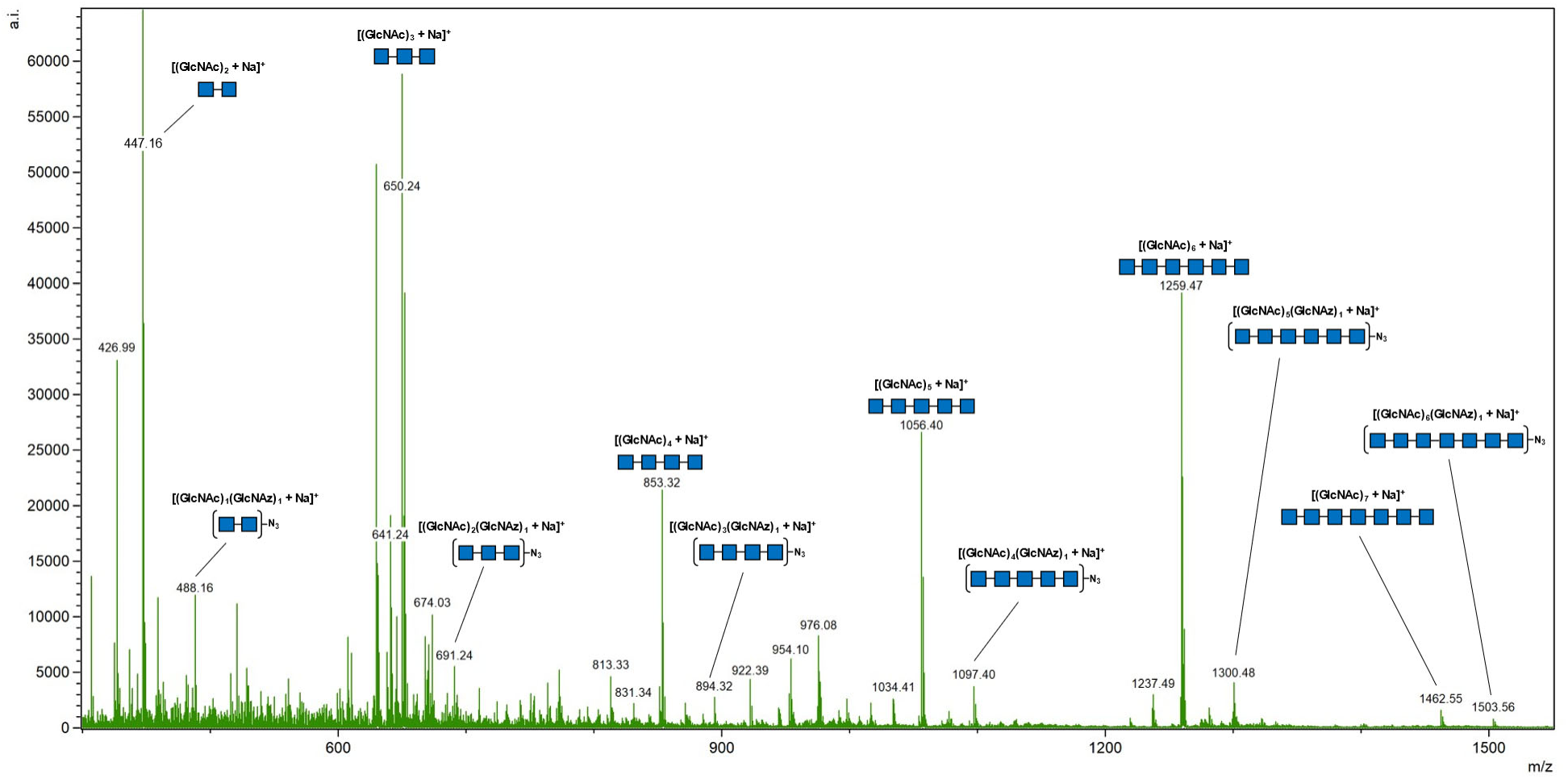
PgaCD reaction mixture using UDP-GlcNAc (20 mM) and UDP-GlcNAz (5 mM) as sugar donors was digested with DspB. Dominant peak of each species is the sodium adduct [M+Na]^+^.

**Figure S23.**
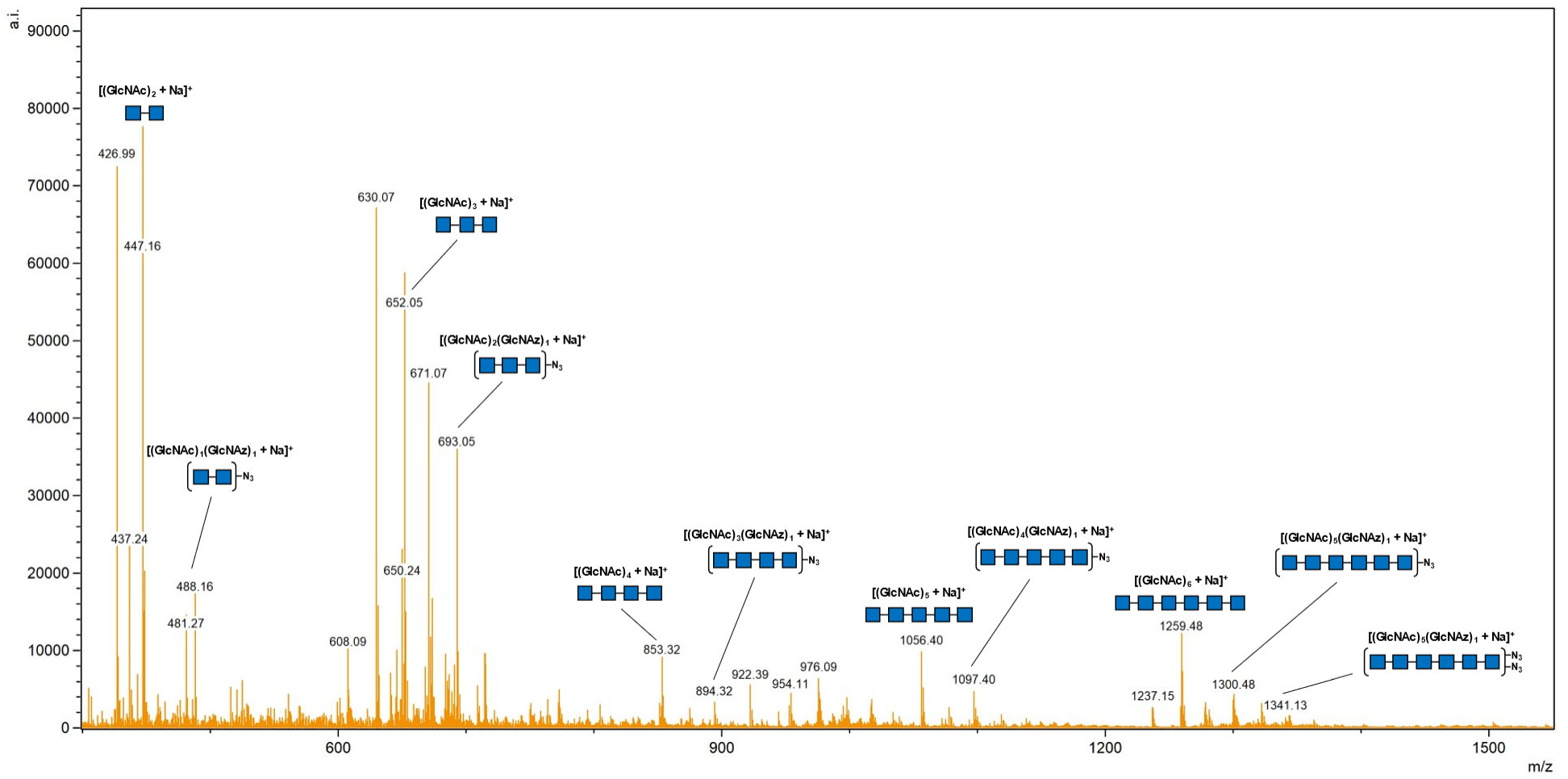
PgaCD reaction mixture using UDP-GlcNAc (12.5 mM) and UDP-GlcNAz (12.5 mM) as sugar donors was digested with DspB. Dominant peak of each species is the sodium adduct [M+Na]^+^.

**Figure S24.**
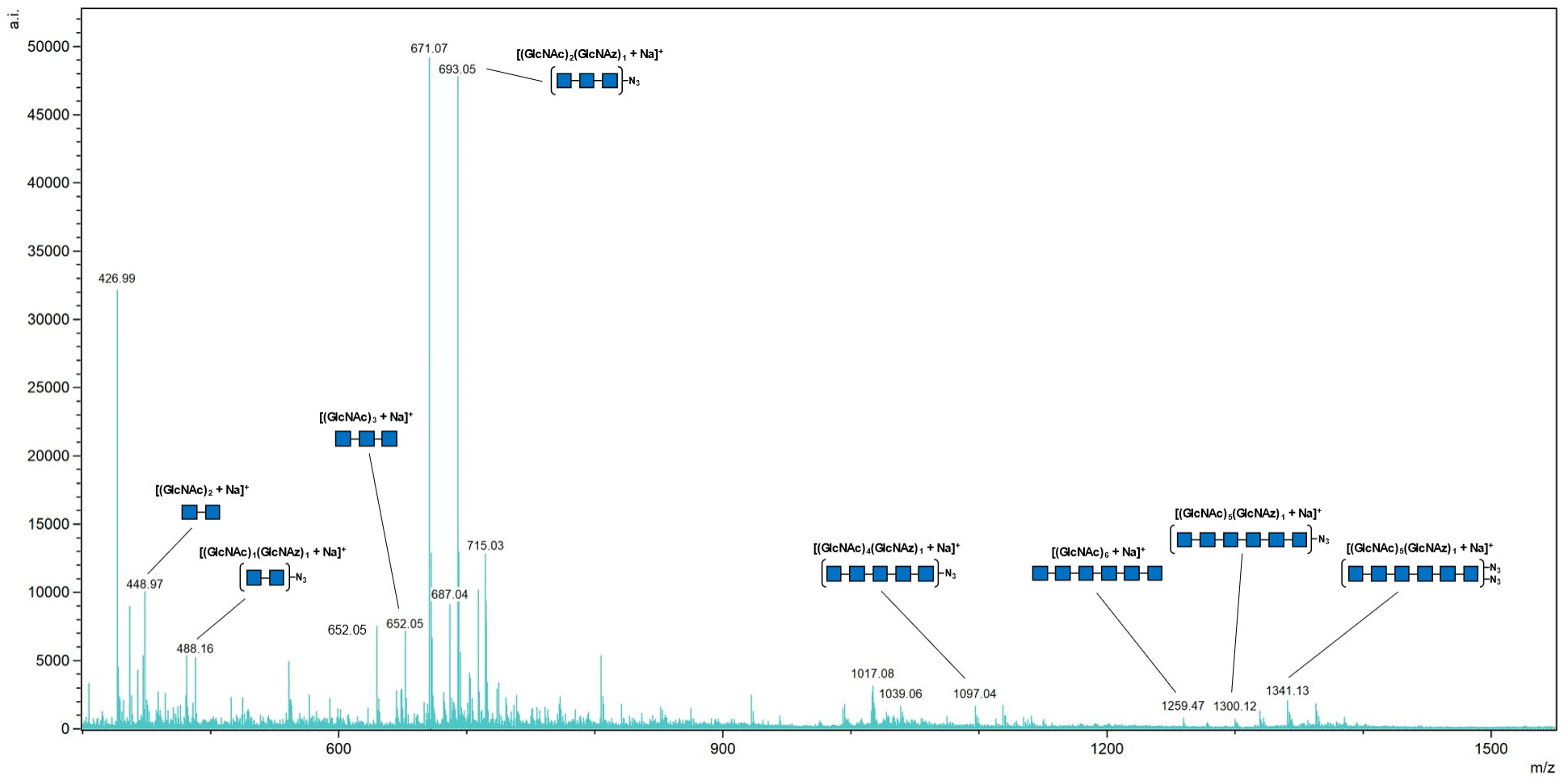
PgaCD reaction mixture using UDP-GlcNAc (5 mM) and UDP-GlcNAz (20 mM) as sugar donors was digested with DspB. Dominant peak of each species is the sodium adduct [M+Na]^+^.

**Figure S25.**
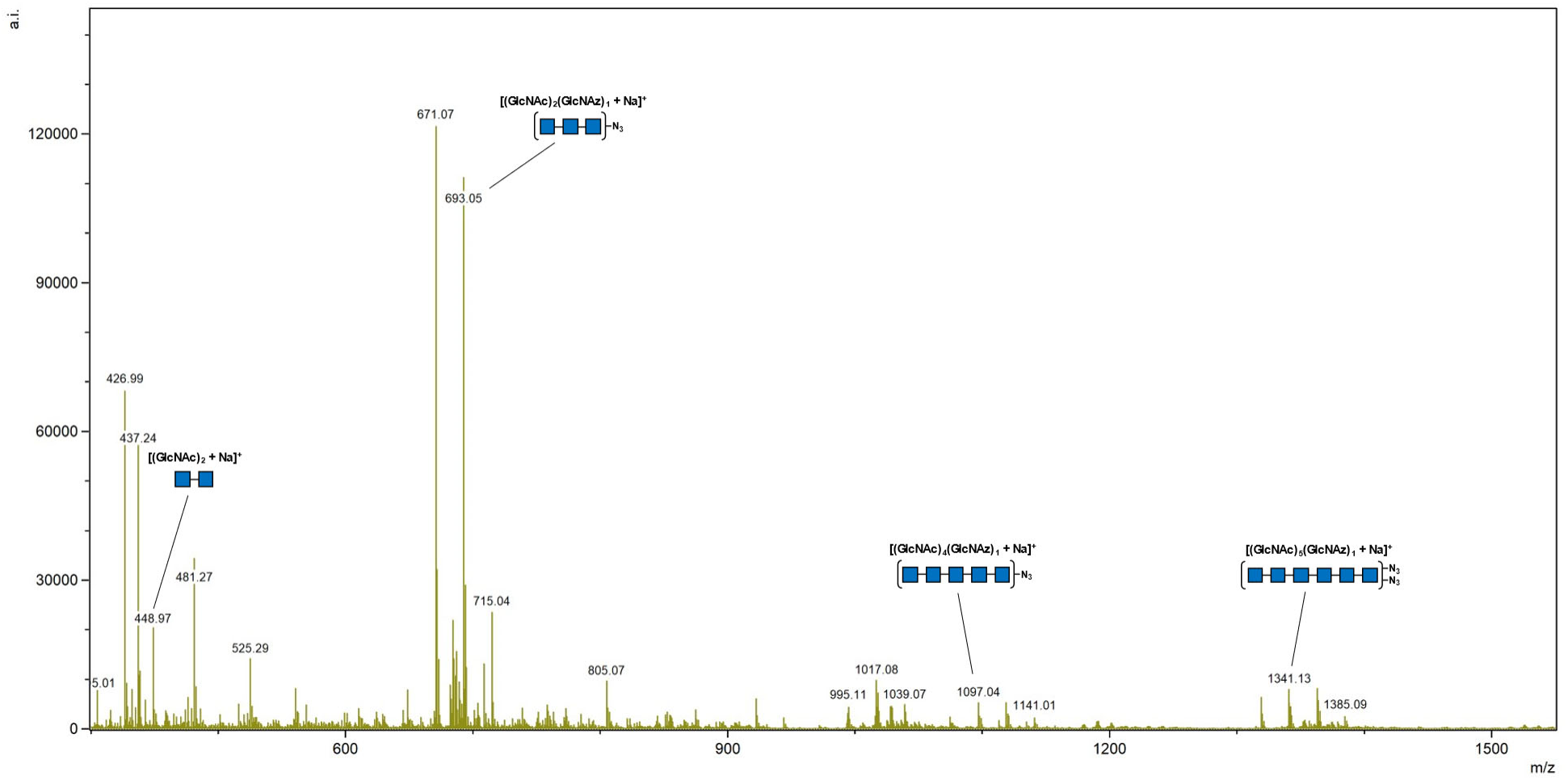
PgaCD reaction mixture using UDP-GlcNAz (25 mM) as sugar donor was digested with DspB. Dominant peak of each species is the sodium adduct [M+Na]^+^. The presence of other non azide-containing GlcNAc species is attributed to low levels of UDP-GlcNAc present in the crude membrane preparations of PgaCD.

